# Genome size, genetic diversity, and phenotypic variability imply the effect of genetic variation instead of ploidy on trait plasticity in the cross-pollinated tree species of mulberry

**DOI:** 10.1101/2023.04.02.535280

**Authors:** Belaghihalli N Gnanesh, Raju Mondal, GS Arunakumar, HB Manojkumar, Pradeep Singh, MR Bhavya, P Sowbhagya, Shreyas M Burji, T Mogili, V Sivaprasad

## Abstract

**Genome size, genetic diversity, and phenotypic vitiation were estimated to develop a model-based population structure and identify ploidy-associated traits of a wide collection of cross-pollinated, highly heterozygous tree species of *Morus*.:** Elucidation of genome size (GS), genetic and phenotypic variation is the fundamental aspect of crop improvement programs. Mulberry is a cross-pollinated, highly heterozygous tree eudicot, and comprised of huge ploidy variation with great adaptability across the world. However, because of inadequate information on GS, ploidy-associated traits, as well as the correlation between genetic and phenotypic variation hinder the further improvement of mulberry. In this present research, a core set of 157 germplasm accessions belonging to eight accepted species of *Morus* including promising functional varieties were chosen to represent the genetic spectrum from the whole germplasm collection. To estimate the GS, accessions were subjected to flow cytometry (FCM) analysis and the result suggested that four different ploidies (2n=2x, 3x, 4x, and 6x) with GS ranging from 0.72±0.005 (S-30) to 2.89±0.015 (*M. serrata*), accounting ∼4.01 fold difference. The predicted polyploidy was further confirmed with metaphase chromosome count. In addition, the genetic variation was estimated by selecting a representative morphologically, diverse population of 82 accessions comprised of all ploidy variations using simple sequence repeats (SSR). The estimated average Polymorphism Information Content (PIC) and expected heterozygosity showed high levels of genetic diversity. Additionally, three populations were identified by the model-based population structure (k=3) with a moderate level of correlation between the populations and different species of mulberry, which imply the effect of genetic variation instead of ploidy on trait plasticity that could be a consequence of the high level of heterozygosity imposed by natural cross-pollination. Further, the correlation between ploidies, especially diploid and triploid with selected phenotypic traits was identified, however, consistency could not be defined with higher ploidy levels (>3x). Moreover, incite gained here can serve as a platform for future omics approaches to the improvement of mulberry traits.

## Introduction

Mulberry (*Morus* spp.) has been commercially exploited as the host of monophagous pest silkworm (*Bombyx mori* L.). It belongs to the Moraceae family comprised of 37 genera with more than 1,100 species (Clement and Weiblen, 2009). The genus *Morus* has over 10 species with more than 1,000 cultivated varieties spanning Asia, Europe, Africa, and the United States (He et al., 2013). Efforts were made to classify *Morus* spp., however, to date, taxonomic nomenclature remains doubtful (Zeng et al., 2015). Besides that, genetics of inheritance are also complicated in *Morus* spp. As an account of the occurrence of a higher level of heterozygosity as well as polyploidy (or whole-genome duplication, WGD), one of the most important evolutionary processes in higher plants (Otto, 2007 and Wood et al., 2009).

Cytology studies in mulberry are more complicated because of extreme variations in chromosome size and number (Datta, 1954). However, since 1920, various researchers attempt to understand meiotic behaviour, because it’s considered a very important aspect of breeding programmes. Mulberry has a wide range of ploidy ranging from haploid (2n=2x=14) to docosaploidy (2n=22x=308) (Yamanouchi et al., 2017), for example, *M. notabilis* was reported as “haploid” in nature (2n= 2x= 14, He et al., 2013), and diploid (2n = 2x = 28) *Morus*s pp. such as *M. alba*, *M. atropurpurea*, *M. bombycis*, *M. indica*, *M. latifolia* and *M. rotundiloba* (Datta, 1954). Majority of triploids (2n = 3x= 42) and tetraploid (2n = 4x = 56) have been identified in *M. laevigata* (Das, 1961). Hexaploid species (2*n* =6*x* = 84), such as *M. serrata* (Basavaiah et al., 1989) and *M. tiliaefolia* (Seki, 1952) are also recognized, and earlier reports also suggest that polyploidy can extend up to docosaploidy (2n = 22x= 308) as in *M. nigra* (Basavaiah et al., 1990; Yamanouchi et al., 2017). The economically important species of mulberry available in India are *M. alba*, *M. indica*, *M. atropurpurea*, *M. nigra*, *M. serrata, M. latifolia*, and *M. laevigata* (Dandin et al., 1987; Basavaiah et al., 1989). Moreover, metaphase chromosome count, detecting the ploidy status, size, and nature of chromosomes and behaviour in pairing and segregation during meiosis has been restricted to a few popular mulberry varieties and polyploid species only like *M. serrata, M. tiliaefolia* and *M. nigra* (Basavaiah et al., 1990; Chakraborti et al., 1999; Thumilan and Dandin, 2009; Venkatesh, and Munirajappa, 2013; Venkatesh et al., 2013; 2014; Venkatesh, 2014; 2015; Shafiei and Basavaiah, 2018). Despite our increasing recognition of the chromosomal number or ploidy level, we are still unaware of the biological effects of ploidy in mulberry. Application of FCM and GS estimations of a large collection of mulberry are missing, which is considered to be a precious way to generate useful information for the analysis of the genetic relationship, phenotypic diversity, estimation of ploidy-associated traits, and other fundamental aspects of mulberry genomes.

In the last few decades, genetic relationship analysis of mulberry is an important component of crop improvement programs and molecular markers are the most popular to understand the diversity between *Morus* species (Awasthi et al., 2004; Vijayan et al., 2022). Primarily, Xiang et al. (1995) applied the Random amplified polymorphic DNA (RAPD) technique to illustrate variation among different species of *Morus*. Later Inter-Simple Sequence Repeats (ISSR) work has been reported on the genetic diversity analysis amongst the mulberry plants grown in India (Awasthi et al., 2004; Zhao et al., 2006). Additionally, several restrictions limit the estimation of the genetic diversity of mulberry as highlighted by earlier reports like-(a) lack of sufficient molecular markers in mulberry (Mathithumilan et al., 2016); and (b) lack of codominant markers to construct the genetic map, which is considered to be a major constraint towards marker-assisted breeding for targeted trait improvement (Pinto et al., 2018; Vijayan et al., 2022). As mulberry has a highly cross-pollinated heterozygous genome, Simple Sequence Repeats (SSRs) are the most appropriate DNA markers to use for analyzing the genetic diversity of mulberries (Zhang et al., 2006; Mathithumilan et al., 2016; Arunakumar et al., 2021; Shinde et al., 2021). SSR-based approaches appear to be one of the best DNA markers for detecting genetic diversity within a species for several reasons, especially for revealing high levels of polymorphisms (Orhan et al., 2020). In *Morus* spp. SSR markers developed from *M. indica* are used to illustrate genetic relationships and origin of cultivated mulberry (Krishnan et al., 2014). However, limited studies on SSR markers-based genetic diversity and model-based population structure analysis of a wide collection of mulberry germplasm have been conducted.

In brief, in the present study, we have tried to highlight a few fundamental aspects of mulberry core set collections like (1) variation of GS (FCM-based), (2) genetic diversity (SSR-based), and (3) phenotypic variation (growth-yield and reproductive trait-based) of a wide collection of cross-pollinated, highly heterozygous tree species of genus *Morus* to develop a model-based population structure and identify ploidy-associated traits. The information generated in the present study may serve as a platform for future omics approaches for the improvement of mulberry traits.

## Materials and Methods

### Plant materials

A total of 157 accessions of the eight different *Morus* species including popular varieties cultivated in India were used in this study, which was obtained from the core collection maintained at the 1). Central Sericultural Germplasm Resources Centre (CSGRC) (csgrc.res.in), Hosur, India; 2). Central Sericultural Research and Training Institute (CSRTI), Mysore, India; and 3). Karnataka State Sericulture Research and Development Institute, Karnataka, India. The common name, accession number, species name, and place of origin (**Table S1; Figure S1**). Accepted nomenclature for the synonymous and unresolved species was considered from The Plant List (theplantlist.org) database (description available in **Table S1**). As per The Plant List (theplantlist.org) database, most of the studied germplasm accessions have belonged to accepted eight species viz. *M. alba* L., *M. auastralis* Poir., *M. cathayana* Hemsl., *M. indica* L., *M. macroura* Miq., *M. nigra* L., *M. rubra* L., and *M. serrata* Roxb. and remaining species like *M. bombycis* Koidz. (Synonym of *M. auastralis*), *M. laevigata* Wall. ex Brandis (synonym of *M. macroura* Miq.), *M. latifolia* Poir. (Synonym of *M. alba* L.), *M. multicaulis* Perri. (Synonym of *M. alba* L.), *M. rotundiloba* Koidz. (Unresolved), and *M. sinensis* Loudon. (unresolved) used in this study were identified and merged with the accepted eight species, to avoid taxonomical/classification-related constraints.

### Genome size estimation using flow cytometry (FCM)

To estimate the genome size, flow cytometry (FCM) analysis was performed by using the modified protocol as described by Galbraith et al., (1983). For analysis, nuclei were isolated from 40 mg of fresh young leaves (∼5 mm^2^) of mulberry along with the internal reference standard *Pisum sativum* (2C = 9.09 pg, diploid) as mentioned by Mondal et al., 2023a. The tissue sample was taken in a Petri dish comprised of 2 ml of buffer solution (Hypotonic propidium iodide, 50 μg/ml in 3 g/L trisodium citrate dihydride containing 0.05% (v/v) of non-ionic detergent equivalent (Igepal CA-630) containing 2 mg/mL RNase A) and immediately co-chopped to homogenate using a sharp razor blade for 60 sec at 4^0^C. The homogenate solution was filtered through a 37 µm nylon filter to remove large debris. Samples were incubated at 37 ^0^C for 30 min and FCM analysis was performed using BD LSRFortessa™ cell analyzer (BD Biosciences). Signals of PI-stained nuclei were processed with an excitation filter and absorbance filter by the flow cytometer. For each accession, three samples were examined and analyzed using BD FACSDiva 8.0.3 Software. Genome size (in megabase pairs, Mbp) was calculated according to the formulae given by Dolezel et al. (2003) [Ploidy= Reference ploidy × (mean position of the G1 sample peak/mean position of the G1 reference peak)] with the conversion of 1 pg equal to 980 Mbp.

### Chromosome number

Besides FCM, metaphase chromosome count was performed to validate FCM data using the protocol as described by Mondal et al. (2023a). In brief, shoot tips of 12 selected accessions (**Table S2**) were collected at 9.30-10.00 AM in 1 ml of saturated *p*-dichloro benzyne (PDB) with 2 drops of 8-hydroxyquinoline (8HQ) in 1.5 ml Eppendorf. After the collection shoot, meristems were transferred to a 0 °C freezer for 5 min and it is subjected to transfer to 4 °C for 16-20h. After that, meristems were transferred to ice-cold 3 parts of 100% ethanol: 1 part of glacial acetic acid and incubated at RT for 1h. Then samples were transferred to 4 °C for 3 days. After that, samples were incubated with the enzymatic solution comprised of pectinase (2.5% w/v), cellulase (1%), and pectolyase (2 %) for 4-6h. Then, samples were transferred to 1% orcine stain for 24h at RT, and during microscopy 2 drops of Acetoorcine stain were used to squash the sample. Chromosome count was performed using Cilika (Madeprime, India) microscope.

### Molecular analysis

Eighty-two germplasm were selected based on collection site, open/cross-pollinated hybrid, and breeding information (functional hybrids, natural hybrids, cultivars, etc.) and different ploidy levels covering all available *Morus* spp (82 accessions of Table S1) for genetic diversity analysis. For DNA isolation, 1-2 younger leaves were selected and the modified CTAB method was followed (Anuradha et al., 2013). The quality of genomic DNA was examined by agarose (0.8%) gel electrophoresis and quantified using a Nanodrop DNA quantifier (Thermo Fisher Scientific, NanoDrop 2000 UV-Vis). Reliable and reproducible twenty polymorphic nSSR primer pairs were selected after the screening of 62 nSSR markers for molecular characterization (Mathithumilan et al., 2016; Pinto et al., 2018, Primers in **Table S3**). The SSR genotyping was carried out in 10 μl of PCR reaction containing ∼50 ng of DNA, 2x PCR Amplicon master mix, and 10 picomoles of SSR primers (Eurofins Pvt. Ltd., Bengaluru) using the PCR program: 94°C for 5 min, followed by 35 cycles of 94°C for 45sec, specific annealing temperature (50°C to 56°C) for 30 sec, 72°C for 45 sec for extension, and72°C for 8 min for the final extension using a thermal cycler (GeneAmp 9700, Applied Biosystems, USA). The final PCR product was separated on 3% agarose gel containing 0.5μgethidium bromide in 1X TBE buffer and gel images were documented in the gel documentation system (Gene Genius, Syngene, UK). Twenty polymorphic SSR marker data were numerically coded as presence = 1, absence = 0, and missing data were coded as ‘-1’ as suggested in the GenAlExV6.5 user manual (Peakall and Smouse, 2012). Gel images of twenty polymorphic SSR markers of 49 representative accessions of mulberry are presented in **Figure S2** along with the scored SSR data (**Table S4**).

### Genetic diversity parameters and Polymorphism Information Content (PIC)

Genetic diversity parameters such as the observed number of alleles (*N_a_*), the effective number of alleles (*N_e_*), observed heterozygosity (*H_o_*), expected heterozygosity (*H_e_*), and Shannon’s information index (*I*) for each SSR marker were computed using POPGENE v. 1.32 (Yeh et al., 1997), while Polymorphism Information Content (PIC) using CERVUS 3.0.7 (Kalinowski et al., 2007). Whereas, GenAlEx V6.5 was used for species-wise diversity measures i.e. percentage of polymorphic loci (%P), *N_a_*, *N_e_*, *I*, *h* (diversity) and *uh* (unbiased diversity), pairwise population matrix of Nei’s genetic distance and pairwise PhiPT values.

### Population structure and Cluster analysis of *Morus* using SSR markers

Mulberry germplasm individual molecular data were scored from SSRs banding patterns and STRUCTURE software (Pritchard et al., 2000) was used to infer the population structure of 82 mulberry accessions. An admixture model was used with the option of correlating allele frequencies between populations. Ten runs were conducted for each value of the number of populations (K), with K ranging from 1 to 12. The length of burn-in Markov Chain Monte Carlo (MCMC) replications was set to 500,000 and data were collected from over 500,000 MCMC replications in each run. We identified the optimal value of K using both the ad hoc procedure described by Pritchard et al. (2000) and the method developed by Evanno et al. (2005) with the help of Structure Harvester software Earl and Von (2012). Mulberry accessions were assigned to the subpopulations based on the cluster assignment probability (Q). Further cluster analysis of 82 germplasm was determined using DARWIN software, version 6.0.21 (Perrier and Collet, 2006). Principle Coordinate Analysis (PCoA) and Analysis of Molecular Variance (AMOVA) were performed using the software GenAlEx V6.5.

### Morphological characters

To correlate phenotypic traits with ploidies, morphologically unique 80 mulberry accessions were chosen to represent the genetic spectrum from the whole germplasm collection. The data was recorded for 9 characters viz. inter-nodal distance (cm), leaf-lamina length (cm), leaf-lamina width (cm), leaf area (sq. cm), petiole length (cm), petiole width (cm), mature inflorescence length (cm), mature fruit length (cm) and mature fruit width (cm) for the two seasons during 2020 and 2021. All the characters were recorded based on DUS test guidelines for Mulberry (<u>plantauthority.gov.in/sites/default/files/mulberry.pdf</u>) and FAO (<u>fao.org/3/AD107E/ad107e0v.htm#TopOfPage</u>). To estimate the inter-nodal distance (InD, cm), the total length of the longest shoot was divided by the total number of nodes after 90 days of pruning. For, leaf-lamina length (LLL, cm) fully grown leaves from the 7^th^ to 9^th^ position in the longest shoot were selected and measured the leaf blade length from the leaf base at the juncture of the petiole attachment to the leaf tip. Additionally, for leaf-lamina width (LLW, cm), the width of the leaf was taken from the widest point on both sides of the leaf margins. The leaf area (LA, cm^2^) of the mulberry was measured using a leaf area meter (Biovis PSM L3000). To measure Petiole length (PL, cm), the petiole portion was separated from the base of the leaf blade and measured, and for petiole width (PW, cm), the thickness was measured using Vernier caliper from the same petioles that were used for measuring PL.

To measure the mature inflorescence length (MIL, cm), fully bloomed inflorescence of male (before anthesis), female (at the receptive stage), and bisexual catkins depending upon the sex expression of the accession were used and measure the length including the pedicel of the inflorescence in a minimum of three inflorescence/plant and three plants/accession. For the measurement of mature fruit length (MFL, cm) fruits fully ripened was collected and the length of the full fruit including the peduncle was recorded, whereas for mature fruit width (MFW, cm) thickness was recorded using Vernier calipers. Parameters like Ind, LLA, LLW, LA, PA, and PW were measured after 90 days of pruning, whereas, MIL, MFL, and MFW were measured after 60 days of pruning.

### Statistical analysis

The normality test (Shapiro-Wilk test) and homogeneity of variance test (Levene’s test) were carried out (the dependent variable for each group is not normally distributed and variances of the groups are not equal). Kruskal-Wallis H test followed by Dunn’s post hoc tests (p < 0.05) was conducted to know the significance among the ploidy groups. All statistical analyses was carried out using IBM SPSS Statistics 23.0. The descriptive statistics (mean, maximum, minimum values, standard deviation, and standard error) were calculated for replicated data. A box plot for ploidy groups was drawn for all characters using the R 3.5.3 version (R Core Team 2018) and R Studio Team (2020). The mean values were used for Spearman rank correlation, ploidy groups PCA, and ploidy-species annotated heatmap analysis with MVappa (https://mvapp.kaust.edu.sa; Julkowska et al., 2019) and ClustVis (https://biit.cs.ut.ee/clustvis, (Metsalu and Vilo, 2015) online tools, respectively.

## Results

### Intra and interspecific ploidy variation of *Morus* spp

The GS estimation of the core collections available in India is not yet investigated so far. Therefore, the estimation of the GS of the 157 potential germplasm encompasses different species of *Morus* that were collected from tropic, subtropical, and temperate areas all over the world, including functional hybrids, popular varieties cultivated in India, and are accounted for in this study (**Table S1; Figure S1**). Results indicate that-

1. GS estimation signifying that the nDNA content of studied mulberry accessions ranged from 0.72 to 2.89pg, the highest amount of 2.89pg DNA was found in *M. serrata* (hexaploid genome) and the least value of 0.72pg was recorded in S-30 (diploid genome) (**Fig. 1A; Table S1**). The nDNA content in the replication of the same accession was almost the same (0.02 to 0.09; **Table S1**).
2. The mean genome content of 0.86, 1.24, and 1.79pg was recorded for diploids, triploids, and tetraploids, respectively. The average GS ranges between 0.71 to 0.98pg for diploids, 1.16 to 1.29pg for triploids, 1.45 to 2.02pg for tetraploids, and 2.88 to 2.91pg for hexaploids are substantially different ploidy levels are differentiated as per the GS estimated (**Fig. 1A**).
3. The mulberry accession in the same species varied slightly (**Fig. 1B**). For example, 0.73pg and 0.98pg of GC were observed in SRDC-2 and Kokuso-20, which belongs to *M. alba*. Similarly, accessions of *M. indica* GS ranged between 0.72pg to 0.98pg in diploids (**Table S1**).
4. Among the predicted four ploidy, accessions of tetraploids showed the highest variability in GS, ranging from 1.45 to 2.02pg (±0.03) compared to diploids and triploids (**Fig. 1A**).
5. Predicted ploidy level indicates wild collections of *M. indica* and *M. alba,* and *M. macroura* comprised all three types of ploidy such as diploid, triploid and tetraploid (**Fig. 1B**). For example, accessions of *M. indica* MI-0173 (1.27±0.031pg), MI-0652 (1.26±0.024pg) and MI-0799 (1.24±0.021pg) are considered to be triploid, whereas MI-0454 (1.69±0.050pg) is tetraploid (**Table S1**). In *M. alba*, accessions MI-0050 (1.16±0.015pg) and ME-0092 (1.21±0.018pg) predicted to be triploid, whereas ME-0149 (1.45±0.024pg) is tetraploid. Accessions of *M. macroura* like MI-0051(1.21±0.018pg), MI-0079(1.23±0.015pg), MI-0521 (1.28±0.035pg) and MI-0772 (1.25±0.025pg) considered to be triploid and MI-0247(1.53±0.048pg), MI-0365(1.91±0.027pg), and MI-0387 (2.02±0.036pg) are tetraploid in nature (**Table S1**).
6. The maximum variability at ploidy levels were observed from *M. macroura,* out of seventeen studied accessions, six are tested as higher ploidy levels (**Fig. 1B**).
7. Additionally, FCM data indicates that with increasing ploidy levels, event count consistently decreased. The less event count might be the consequence of polyploidy-mediated increasing cell size with decreasing cell number (**Fig. 2A-E**).

**Figure 1.**
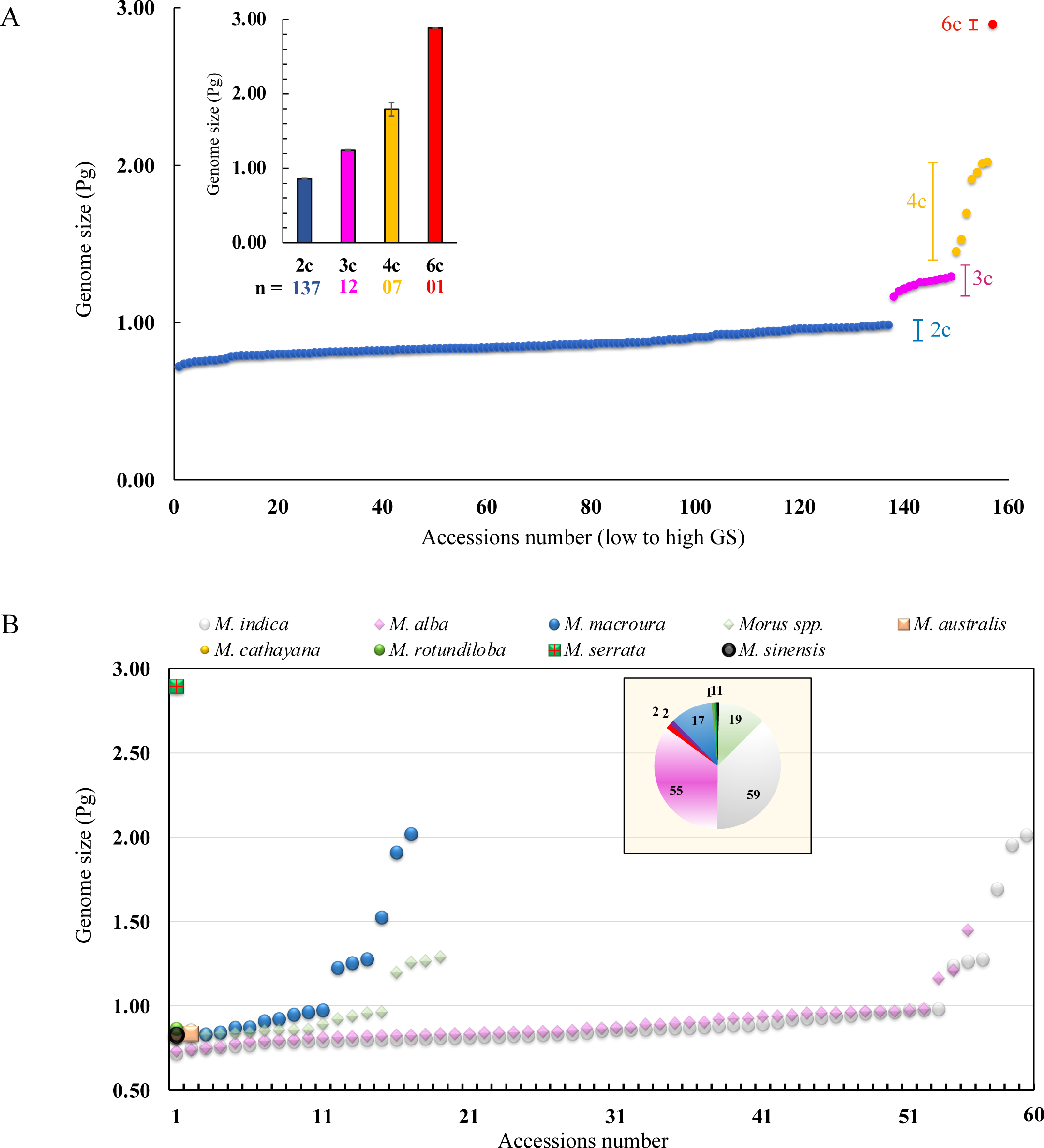
Patterns of genome size (GS) variation in mulberry accession A) Distribution of individual GS estimates and their relationship with ploidy, B) average distribution of GS belonging to diverse *Morus* spp.

**Figure 2.**
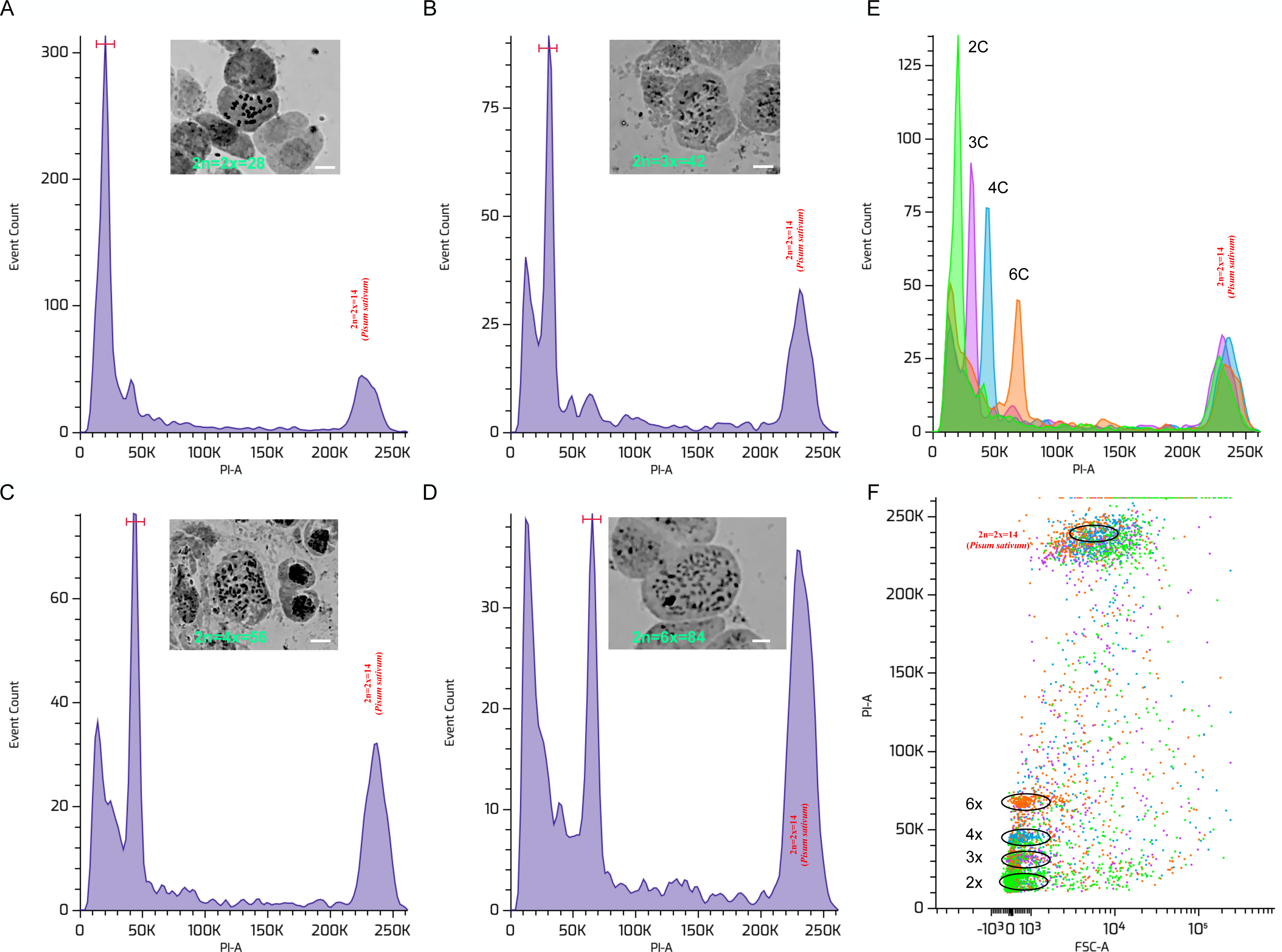
Ploidy detection by flow cytometry and chromosomal counting. Flow cytometry histograms of nuclei extracted from young leaf tissue of mulberry accessions comprising different ploidy groups and the corresponding chromosome numbers such as (A) diploid (2n=2x=28) (B) triploid (2n=3x=42), (C) tetraploid (2n=4x=56) and (D) hexaploid (2n=6x=84) with flow cytometry histograms depicting the relative fluorescence intensity obtained from the simultaneous analysis of isolated nuclei of mulberry leaves and the reference standard (*Pisum sativum*). Distinguishable predicted ploidy levels presented as histogram (E) and scattered plot (F). Average CV for the standards = 3.9 % and for the sample=2.9 %.

### Chromosome counting and validation of ploidy

The present cytological study validated and confirmed the chromosomal numbers of 12 selected mulberry accessions comprising different ploidy groups such as diploid *var*. V1, K2 (2n=2x=28, 415.65 Mbp, 419.08 Mbp), triploid Saranath-1 (2n=3x=42; 612.88 Mbp), tetraploid Lava forest-1 (2n=4x=56; 987.78 Mbp) and hexaploid thick leaf *M. serrata* (2n=6x=84; 1414.84 Mbp) (**Fig. 2A-E**).

### Genetic diversity parameters, Polymorphism Information Content (PIC) and AMOVA

A set of 20 (32.25%) polymorphic SSRs out of 62 SSR primer pairs was expedited to measure the genetic diversity inferences among the 82 mulberry accessions selected based on collection site, open/cross-pollinated hybrid, and breeding information (lines including functional hybrid, culverts, polyploidy, and mutation breeding, etc.). These 20 informative markers generated 94 SSR alleles with an average of 4.7 alleles per SSR marker ranging from 2 (M2SSR112A) to 7 (MULSSR258, MULSSR85, and M2SSR20). A mean Shannon’s Information index (I) value of 1.137 was observed for all the markers varying from 0.383 to 1.914 (**Table 1**). The value of Ho for SSR markers ranging from 0.024 to 0.890 with a mean of 0.477 was observed, while the value of He varied from 0.208 to 0.853 with a mean of 0.606. The average PIC value for SSRs was 0.556. Fifteen (75%) SSR markers in total were found to be highly informative with a PIC value ≥0.50, Four (20%) were moderately informative with PICs values ≥0.25 and <0.50 and the remaining one was least informative with a PIC value <0.25 (**Table 1**).

**Table 1.** Characteristics of the 20 SSR loci among the 82 mulberry accessions

| Locus name | Na | Ne | Ho | He | I | PIC |
| --- | --- | --- | --- | --- | --- | --- |
| M2SSR81 | 3 | 1.422 | 0.085 | 0.299 | 0.525 | 0.273 |
| Moso340-2 | 6 | 3.843 | 0.5 | 0.744 | 1.507 | 0.704 |
| M2SSR87 | 3 | 2.828 | 0.524 | 0.65 | 1.069 | 0.58 |
| Moso288 | 4 | 1.997 | 0.634 | 0.502 | 0.814 | 0.42 |
| MULSSR 253 | 4 | 2.912 | 0.695 | 0.661 | 1.137 | 0.592 |
| M2SSR112A | 2 | 1.49 | 0.39 | 0.331 | 0.51 | 0.285 |
| MULSSR26 | 4 | 3.003 | 0.634 | 0.671 | 1.196 | 0.605 |
| M2SSR68 | 3 | 1.701 | 0.366 | 0.415 | 0.739 | 0.373 |
| MULSSR313 | 5 | 3.242 | 0.598 | 0.696 | 1.303 | 0.65 |
| MULSSR258 | 7 | 4.018 | 0.89 | 0.756 | 1.565 | 0.714 |
| MoSo-157-2 | 6 | 3.335 | 0.293 | 0.704 | 1.331 | 0.652 |
| M2SSR36 | 5 | 2.561 | 0.159 | 0.613 | 1.101 | 0.549 |
| M2SSR1 | 5 | 3.114 | 0.134 | 0.683 | 1.263 | 0.627 |
| MULSSR85 | 7 | 6.583 | 0.768 | 0.853 | 1.914 | 0.831 |
| MULSSR96B | 5 | 3.63 | 0.695 | 0.729 | 1.44 | 0.689 |
| M2SSR10 | 4 | 2.63 | 0.753 | 0.624 | 1.09 | 0.544 |
| M2SSR89A | 5 | 2.465 | 0.39 | 0.598 | 1.107 | 0.532 |
| M2SSR107 | 3 | 1.26 | 0.024 | 0.208 | 0.383 | 0.198 |
| M2SSR82 | 6 | 4.071 | 0.325 | 0.759 | 1.521 | 0.719 |
| M2SSR20 | 7 | 2.596 | 0.675 | 0.619 | 1.226 | 0.577 |
| Average | 4.7 | 2.935 | 0.477 | 0.606 | 1.137 | 0.556 |

Na = observed number of alleles, Ne = effective number of alleles, Ho = observed heterozygosity, He = expected heterozygosity, I = Shannon's information index and PIC = polymorphism information content

The species-wise diversity analysis revealed that in four species a mean of 83.78 % of loci were polymorphic. *M. alba* had maximum polymorphic loci (88.30 %), while *M. indica* had at least one (80.85%). The maximum Na was observed in *M. alba* (1.777) with a mean of 1.686, while Ne (1.487) and I (0.425) were observed in *Morus* spp. with a mean of 1.462 and 0.413, respectively. The h (diversity) ranged from 0. 257 (*M. indica*) to 0.283 (*Morus* spp.) with an average of 0.273 whereas maximum uh (0.302) was recorded in *Morus* spp. with a mean of 0.288 (**Table S5**).

To evaluate the overall partitioning of genetic diversity among studied germplasm collections, an AMOVA was executed using the genetic distance matrix. The AMOVA revealed that 94% of molecular variance was partitioned within species and only 6 % among species (**Table S6**). The overall PhiPT was 0.058 (*p*-value=0.001) suggesting a low level of genetic differentiation among studied *Morus* species. The pairwise population PhiPT and Nei’s genetic distance were analyzed to determine the genetic relationship among *Morus* species (**Table S7**). The highest PhiPT (0.120) and Nei’s genetic distance (0.085) were observed between *M. macroura* and *Morus* spp. revealing that these species are more divergent. Whereas the lowest PhiPT(0.025) and Nei’s genetic distance (0.025) were observed between *M. alba* and *M. indica*, followed by *M. alba* and *M. macroura* with PhiPT(0.039) and Nei’s genetic distance (0.037). These results suggest that the pair of *M. alba* and *M. indica* and *M. alba* and *M. macroura* are genetically closer to each other as compared to other species (**Table S7**).

### Population genetic structure and cluster analysis

The SSR-based clustering analysis of different *Morus* spp. was carried out using multidimensional Principal Coordinates Analysis (PCoA), UPGMA-based Neighbor-joining clustering, and Bayesian model-based complementary clustering approaches. The PCoA (**Fig. 3A**) revealed that *Morus* species were grouped into three major genetic clusters with a high level of intermixing as complemented by UPGMA-based NJ clustering. The first three coordinates of PCoA only explained 25.68% of the total variation (PC1: 9.90%; PC2: 8.94% and PC3: 6.84%; **Fig. 3A**).

**Figure 3.**
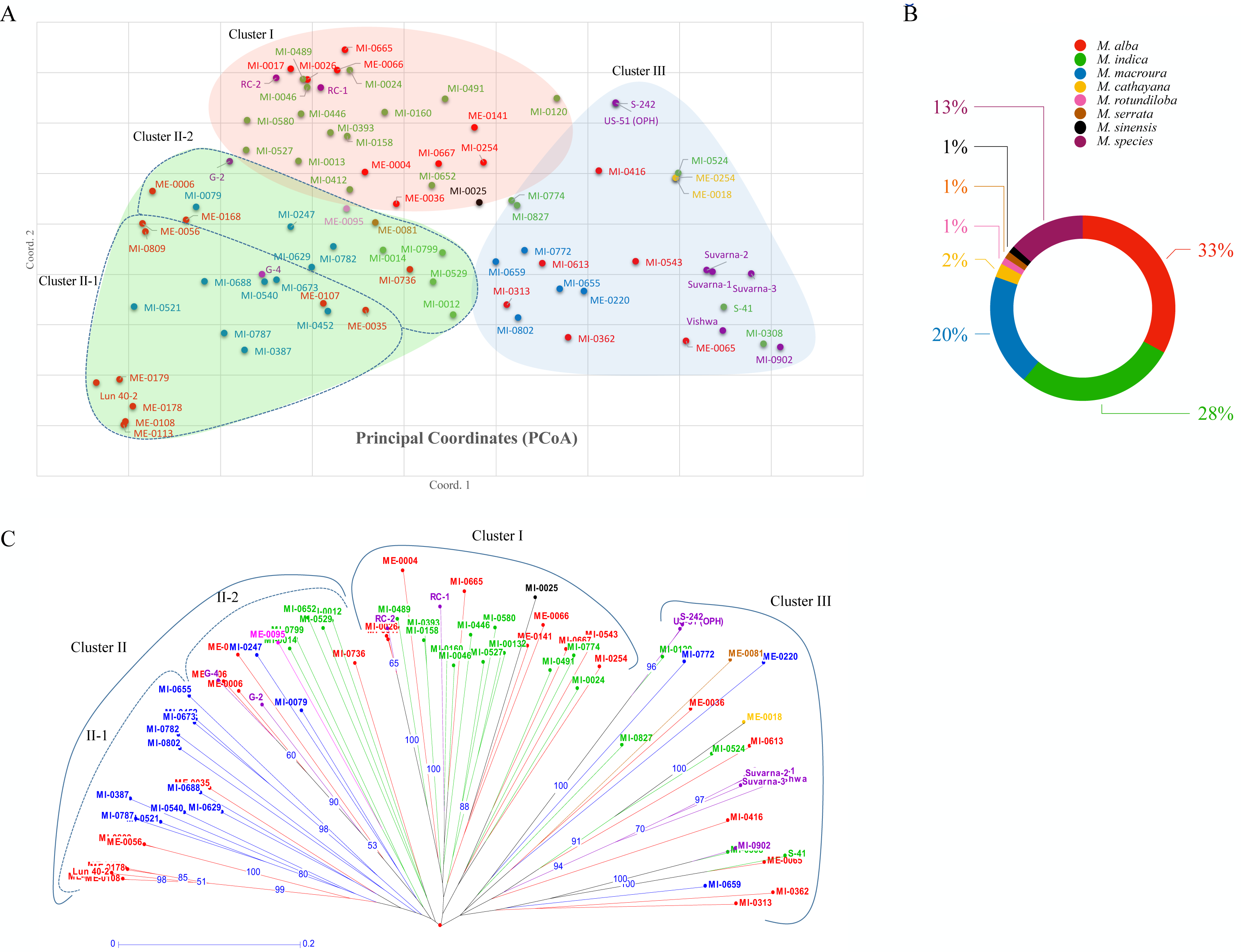
(A) Principal Coordinate Analysis (PCoA) plot of 82 mulberry accessions using SSR markers. The PCoA revealed that 8 *Morus* species were grouped into three major genetic clusters with a high level of intermixing. The first three coordinates of PCoA explained the 25.68% of the total variation (PC1: 9.90%; PC2: 8.94% and PC3: 6.84%). (B) The donut chart represents the percentage of species used in the present analysis. (C) Phylogenetic analysis of 82 mulberry accessions from diverse geographical regions belonging to different species was grouped into three clusters. Different species were highlighted with different colour as described in Figure 5B.

The highest percentage of *Morus* species representation was from *M. alba* (33%), *M. indica* (28%) and *M. macroura* (20%) respectively (**Fig. 3B**). The triploids (Suvarna-1, Suvarna-2, Suvarna-3 and Vishala (MI-0902) were grouped in Cluster III. These triploids were found very close to accession Vishwa, which is one of the parent used in breeding these triploids. Similarly, two accessions of ME-0018 and ME-0254 belonged to *M. cathayana* were grouped in Cluster III and exotic accessions ME-0113 and ME-0108 (Japan), ME-0178 (France), ME-0179 and Lun 40-2 (China) all belonged to *M. alba* and were groped in Cluster II-1 (**Fig 3C**).

Additionally, the UPGMA cluster analysis was used to describe the genetic relationships among the 82 Morus accessions (**Fig. 3C**). According to the dendrogram analysis and dissimilarity coefficient measured using the Neighbor-joining method, it is depicted that the markers were successful in segregating among the mulberry accessions irrespective of species. The dendrogram grouped all accessions into three major genetic clusters-I, II, and III (**See more detailed in Table S8**). Cluster I consists of 25 accessions and the majority of them were *M. indica* (13) and *M. alba* (9) conferring their close genetic relationships as depicted by pairwise population PhiPTand Nei’s genetic distance. Cluster-II consists of 33 accessions and has two sub-clusters, Cluster II-1 and II-2 comprising 19 and 14 accessions respectively. Interestingly, sub-cluster II-1 comprises *M. macroura* (11) and *M. alba* (8) accessions conferring their close genetic relationships. Whereas, the Cluster-III consists of 24 accessions of six mulberry species i.e. *Morus* spp. (7), *M. alba* (6), *M. indica* (5), *M. macroura* (3), *M. cathayana* (2), and *M. serrata* (1) and found to be the highest admixture cluster, since more accessions of which are mulberry varieties developed from two different species (**Fig 3C**). Further, other than cluster I, higher ploidy level accessions are distributed in the entire groups projecting that polymorphic SSRs detected in our study were not able to discriminate among diploids, triploids, and tetraploids.

The existence of population structure in *Morus* accessions was resolute using STRUCTURE analysis based on the determination of optimal K-value (**Fig 4A-C**). These results indicated that *Morus* accessions were grouped into three clusters (**Fig 4C**). The cluster-I represents 35 accessions and the majority of them belong to *M. indica* (16) and *M. alba* (11), cluster-II represents 17 accessions majority of them belong to *M. macroura* (8) and *M. alba* (8), while cluster III represents 17 accessions majority of them belongs to *M. indica* (7), *Morus* spp. (7) and *M. alba* (8). Further, based on membership coefficients (q ≥ 0.70) only 40 (48.78%) accessions were assigned to a specific population structure cluster belonging to pure ancestry. While the remaining 51.22 % of accessions represented a high level of genetic admixtures in three clusters with membership coefficients of ˂ 0.70. Among species, two accessions of *M. cathayana* (100%), eight *Morus* spp. (72.73%), fourteen *M. indica* (60.87%), twelve *M. alba* (44.44%), and four *M. macroura* (25%) accessions represent pure ancestry.

**Figure 4.**
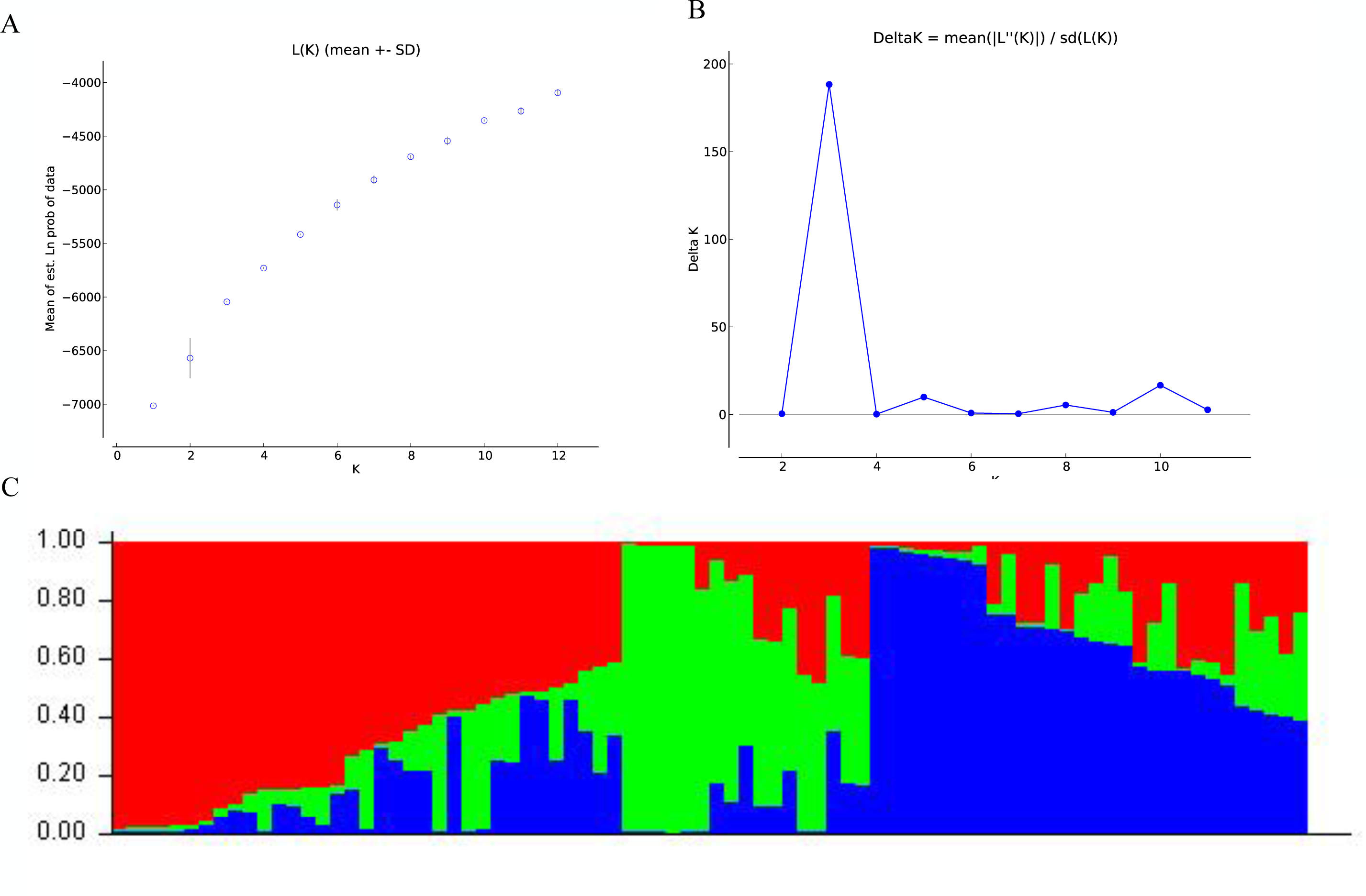
Population structure of 82 mulberry accessions based on SSR markers. A) Log probability of data, L (K) averaged over the replicates, B) Delta K values plotted as the number of subpopulations. (C) Subpopulations, K = 3 inferred using structure analysis.

### Morphological traits variation and correlation

Variations in morphological characteristics among diploids, triploids, tetraploids, and hexaploids were examined and analysed (**Table S9**). From this study, we observed that-

1. The internodal length (InD), leaf-lamina length (LLL), leaf area (LA), and mature inflorescence length (MIL) were greater in triploids compared to diploids and tetraploids (**Fig. 5**, **Table S10**).
2. Leaf-lamina width (LLW) and petiole width (PW) had higher mean values in tetraploids (17.8±6.62 and 0.36±0.1) compared to diploids (13.5±3.57, 0.27±0.09) and triploids (15.9±2.4, 0.32±0.07).
3. Petiole length (PL) was greater in diploids (4.3±1.01) compared to triploids (4.28±0.81) and tetraploids (4.18±1.34). Hence, PL was found to be decreased as the ploidy level increased.
4. Mature fruit length (MFL) was greater in triploids (3.85±0.77) than in diploids (2.42±0.65).
5. However, InD and MFW were significantly higher in triploids (7.83±1.15; 0.97±0.09) than in diploids (5.95±1.74; 0.96±0.19) and mature inflorescence length was significantly higher in triploids (4.05±1.02) than in diploid (2.2±0.84) and tetraploids (2.27±1.38) as per the statistical analysis by Kruskal-Wallis H-test (**Table S10**).

**Figure 5.**
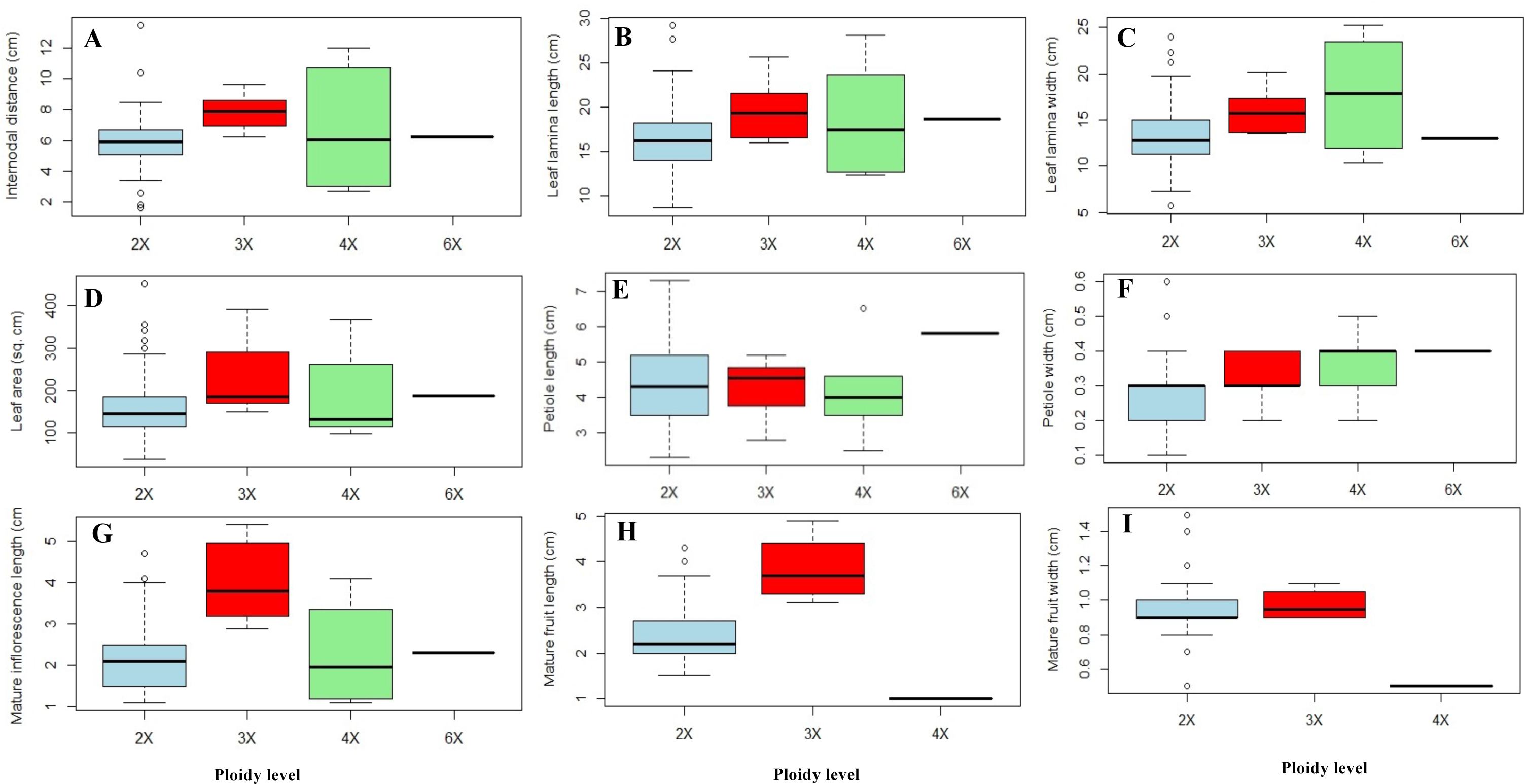
Box plot indicating variation of nine phenotypic traits, A-Inter-nodal distance (cm), B-Leaf-lamina length (cm), C-Leaf-lamina width (cm), D-Leaf area (sq.cm), E-Petiole length (cm), F-Petiole width (cm), G-Mature inflorescence length (cm), H-Mature fruit length (cm) I-Mature fruit width (cm) in the different ploidy levels detected in selected mulberry accessions. Outliers are shown as circle.

The correlation results revealed a significant positive correlation of GS with traits viz., mature inflorescence length (MI, r=0.45), and fruit length (FL, r=0.38) **(Fig. 6A-B; Table S11)**. However, it had recorded a non-significant association with internodal distance (InD, r=0.19). It had a negative association with petiole length (PL, r=-0.04), leaf-lamina length (LLL, r=-0.08), and leaf area (LA, r=-0.02) which is also depicted in **(Table S11)**.

**Figure 6.**
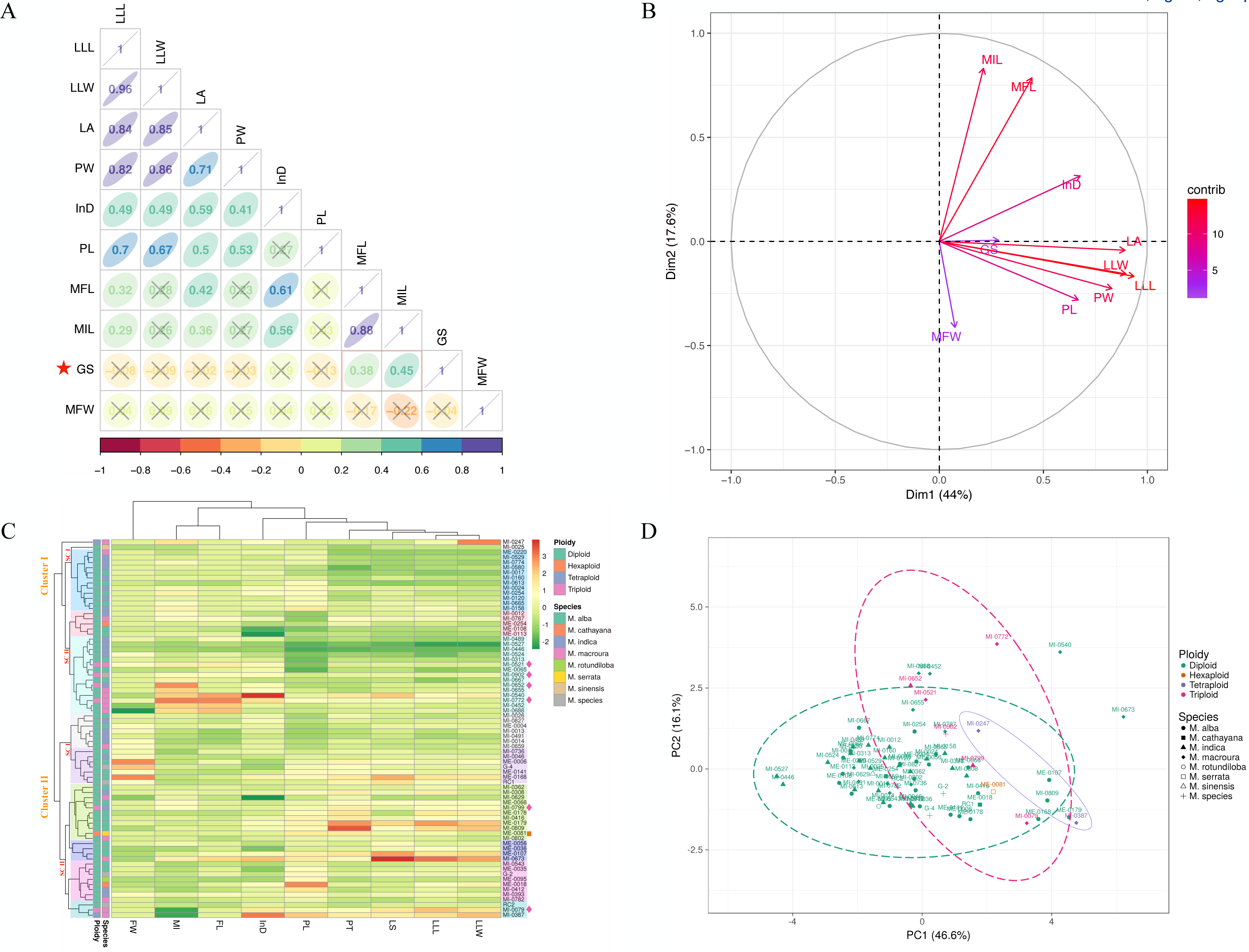
(A) Correlation plots of nine morphological traits and genome size (B) Principal component analysis (PCA) indicates the contribution and relationship between 9 traits to phenotypic diversity and genome size (GS). Colour intensity was used for scaling (0-15 range). (C) Heatmap illustration of clustering relationship between morphological traits with ploidy and species annotation. Colour variation represents the level of trait variation across the species, ploidy as well as accession level. (D) PCA result indicates ploidy-grouping and distribution of studied accessions at the species level. The abbreviation denoted in brackets: leaf-lamina length (LLL), leaf-lamina width (LLW), leaf area (LA), petiole width (PW), inter-nodal distance (InD), petiole length (PL), mature fruit length (MFL), mature inflorescence length (MIL), genome size (GS) and mature fruit width (MFW)

Ploidy-species annotated phenotypic data was represented in hierarchical clustering with heat maps and principal component analysis (**Fig. 6C,D**). Results indicate that 82 accessions were clustered into Cluster I and Cluster II, whereas, each cluster was divided into two sub-clusters (SCI and SCII). Primary groupings for entries were denoted by different colours (9 groups). Cluster I comprised one tetraploid, four triploids, and twenty-nine diploids. On the other hand, Cluster II have comprised of hexaploidy, one tetraploid, two triploids, and thirty-six haploids. Hence, out of six, four triploids were grouped in Cluster I (SCII) and higher MIL and MFL were observed in this cluster of genotypes. In general, the heat map demonstrated that Cluster I comprised triploids have higher MIL and MFL than diploids. Although, the value of LLW, LLL, LA, PT, and PL were higher for the genotype belonging to diploids as compared to other ploidies. Additionally, PCA data (**Fig. 6 C,D**) indicates that the expansion of traits was restricted for diploids as compared to triploids. Data also suggests that few accessions of diploids have a higher value for specific traits. For example, accessions MI-0540 and MI-0673 were outgroup 95% confidence circles because of a higher value of inter-nodal distance (InD, 13.50 cm) and leaf area (LA, 451 cm2), respectively.

## Discussion

### Estimation of genome size (GS) and intra-specific variation

Germplasm is an enduring resource management mission and asset for civilization and to fully explore the potential of germplasm material, particularly for the complex traits associated with yield in the face of the current threats of climate change, it is necessary to reconcile research advancement and innovation of practices (Mondal et al., 2023b). There are several crop germplasm characterized in reference to ploidy level variation, and genetic and phenotypic diversity to explore the potential of germplasm material as well as elucidated consequences of polyploidization on functional traits (Liang et al., 2019; Zhang et al., 2019; Guo et al., 2022; Kumar et al., 2022; Samarina et al., 2022). Additionally, polyploidy (or whole genome duplication, WGD) has been recognized as a source of evolutionary force and its role in species diversification has been well understood (Van de Peer et al., 2021). Despite its importance, genome size (GS) variation across the species and ploidy-level have not been considered in germplasm material like tree plants mulberry, therefore it remains to be explored for a better understanding of fundamental aspects like (1) expansion of polyploidization/WGD occurred in mulberry; (2) preferred polyploid advantages present among the ploidy groups; (3) specific leaf/fruit traits associated with polyploid; and (4) how is population structure affected by polyploidization or WGD.

In the present section, we discuss the status of available polyploids of mulberry to understand polyploidy variation across the species level and try to address the first question. To address how and to what extent WGD occurred in mulberry, 157 mulberry accessions belonging to seven different species including popular varieties cultivated in India were selected and subjected to genome content analysis using FCM. Genome content (GC) data indicates that it ranged from 0.72 (*M. indica*, diploid) to 2.89 pg (*M. serrata*, hexaploid). Thus, the smallest genome size: highest genome size content ratio is about 4.01 (2.89/0.72) was observed within the studied population. Previous genome content studies revealed the 2C value of mulberry species ranged between 0.70 to 0.73 pg for diploid, 1.04 to 1.17 pg for triploid, 1.42 to 1.63 pg for tetraploid, and 2.02 to 2.48 for hexaploid, and 7.26 pg for docosaploid (Yamanouchi et al., 2017). Thus, the present results corroborate with a previous study that a higher level of variation is present both at the species level as well as within the accessions of the same genus. However, the estimated GS of diploid *M. alba* differed from the previous study by Ohri and Kumar (1986), it was around 1.7pg for the n=14. Whereas, diploid *M. alba* was estimated to be 1.7 pg/2C measured by micro densitometry using Feulgen stained which might have some technical shortcomings (Greilhuber, 1998, Chang et al., 2018).

In our study the intraspecific variations of diploid GC data indices that about 34%, 37%, and 18% variation among the genotypes belonging to *M. indica* (59 genotypes), *M. alba* (55 genotypes) and *M. macroura* (17 genotypes), respectively. Moreover, GC variations in terms of the fold change is about 2.79, 1.98 and 2.43 among diploids belonging to *M. indica*, *M. alba* and *M. macroura*. Intraspecific variation has been recorded in *Morus* (Chang et al., 2018) and suggested the GS variation occurred by detection method, or by environmental factors, such as water stress, soil nutrition, or extreme climate condition, and with genetic factors, including duplications, or loss of some non-essential genes in an ongoing evolutionary process that increased or decreased the genome size (Benett and Leitch, 1995; Chang et al., 2018, Greilhuber, 1998; Greilhuber and Leitch, 2013; Laurie and Bennett, 1985; Rayburn et al., 1985). Recently, Xuan et al., (2022) illustrated that chromosome fissions/fusions were the key mechanisms underlying the evolution of differences in the basic chromosome number between *M. notabilis* and *M. alba*.

Variation in terms of genotypes from other species like *M. rotundiloba*, *M. cathayana*, *M. australis*, *M. serrata*, *M. sinensis* is non-significant because of their representation in small numbers (1-2 genotypes). Hence, for diploids, more variation in genome content values was observed in *M. indica* and *M. alba*. Meanwhile, *M. macroura* showed a high level of variation in tetraploids as previously observed by Yamanouchi et al., (2017). However, these estimates differ from previous FCM assessments, possibly because of differences in the DNA reference standard, dyeing time, and fluorochrome difference in the current study (Doležel et al., 1992; Guo et al., 2015). Propidium iodide (PI) is commonly referred to for GS estimation through FCM as used in our present study (Doležel et al., 1992, Yamanouchi et al., 2017).

That GS variability does occur among the various genotypes at an intraspecific level due to the involvement of different genomes during hybridization and selection as well as naturally. Further GS variations are also observed among the genotypes collected from different ecological conditions. Mulberry is reported to be a higher adaptability tree plant with a huge variety of species and ploidy. The use of wild species in breeding provides a lot of possibilities, to incorporate the traits such as drought, frost, and disease resistance into cultivated varieties (Tikader and Kamble, 2008). Breeding of mulberry takes about 12-16 years and to develop a variety, ploidy compatibility is an essential aspect of breeding. To date, the varieties (like V1, K2, S13, MSG2, etc.) released by the researchers are not sufficient for the future sericulture industry, and an urgent need to develop climate-resilient mulberry var, including diploids and triploids. Ploidy and GS can influence reproductive compatibility, fertility, and heritability of traits hence our data can be useful for understanding the breeding potential between the different species of mulberry. In this context, we have provided a new platform for future ploidy breeding by focusing on GS and diversity analysis of mulberry accessions available in India and their relationship.

Besides the FCM analysis, classical cytological work to estimate the metaphase chromosome number is essential to validate the chromosomal number, which is lacking in previous studies (Ohri and Kumar, 1986; Yamanouchi et al., 2017; Chang et al., 2018). Attempts were made to count metaphase chromosome numbers using shoot-tips of original plants and present cytological evidence corroborates with previous studies (Basavaiah et al. 1989; Venkatesh, 2015; Venkatesh Munirajappa, 2013; Venkatesh et al. 2014; Kumara and Basavaiah, 2016; Yamanouchi et al 2017; Kumara et al. 2021), which strengthening our FCM analysis.

Till 2020, chromosome configuration, genome size, meiotic association, and chromosome assembly at the genetic level were restricted in *Morus*. However, the controversy over the ploidy of *M. alba* was clarified by Jiao et al., (2020) that the chromosome number (2n) of *M. alba*, is 28 (diploid) which provides a basis for a better understanding of the developmental and regulatory mechanism of economic target traits. Moreover, present GS estimation for diploids, triploids, tetraploids, and hexaploids, across the species level, was found to be technically foolproof to discriminate the higher ploidy level in the mulberry, because, among the different methods for GS estimating, FCM is now considered as the most precise method (Pellicer and Leitch 2014; Bourge et al., 2018). GS varies significantly across plant life; however, variation of GS in vascular plants is associated with evolutionary consequences on plant development and ecological performance (Vesely et al., 2012; Greilhuber and Leitch, 2013). Moreover, the present research generated GS of 157 core collections, in turn, the information can be useful for effective genetic improvement and genetic diversity investigations. Additionally, the present result and discussion clarify that *Morus* spp. having a higher level of GS variation, not restricted at the species level but also within the accessions of the same genus.

### Genetic Diversity Based on SSR Markers

Understanding of genetic diversity of mulberry is another important aspect of the future breeding program. Genomics-based breeding was reported to be an effective and promising approach to reaching sustainable crop improvement and which is more relevant specifically for perennial crops such as mulberry (Mathithumilan et al., 2013). Despite the significant progress achieved through conventional breeding, even though to date it is distressingly slow, mainly because of the perennial growth habit and complex inheritance pattern (Mathithumilan et al., 2013). Additionally, the adoption of modern genomic approaches for crop improvement is severely constrained by the lack of sufficient molecular markers in mulberry (Mathithumilan et al., 2016). Finally, the paucity of genomic information for co-dominant marker systems has been a serious obstacle to using molecular breeding for the advancement of mulberry genetics (Pinto et al., 2018).

SSRs are simple tandemly repeated DNA sequences comprising units of 01 to 06 nucleotide(s) and are ubiquitous in plant genomes (Tautz and Renz, 1984; Zhong et al., 2021). Because of their distinct qualities of co-dominant inheritance, multi-allelic nature, vast genome coverage, high abundance, and especially good reproducibility, SSRs are the most widely recognized genetic markers that are actively used in plant breeding (Powell et al., 1996). SSR markers are extensively employed in plant research for a wide range of studies including DNA fingerprinting, high-density mapping, genome comparative mapping, population diversity study, and assisted breeding (Zhong et al., 2021). Although the genetic distances, evolution, and kinship of *Morus* have been studied by using molecular marker technology, the conclusions are still controversial (Zhao, 2005; Muhonja et al., 2019). In the present research, a set of stable and highly polymorphic SSR markers were employed and the average polymorphism information content (PIC) value for SSRs was 0.556 and the observed heterozygosity was 0.477. Hence, markers’ inheritance reflected together with relatively high polymorphic information content (PIC) with an observed heterozygosity value of 0.477, clearly explaining their potential as genetic resources in diversification study. Moreover, information generated in the present research based on SSR marker-based genotyping, higher PIC and observed heterozygosity can be efficiently used for future molecular breeding programs of the mulberry species/genotypes. Recently, Jain et al., (2022) reported single-nucleotide polymorphism (SNPs) ranging from 241,897 (S1) to 657,137 (*M. rotundiloba*) in different mulberry accessions such as K2 reference genome with other accessions namely Punjab Local, *M. indica,* BR-8, *M. multicaulis,* and *M. serrata* showed higher genetic variations with ≥0.5 million SNPs which is conformity in our finding that phylogenetic analysis through SSRs revealed diverse relationship between selected species of *Morus* genome. SSR analysis of the mulberry germplasm confirmed a complicated genetic background and high genetic diversity. However, SSR markers were not able to differentiate among ploidy levels hence flow cytometry method is reliable for the determination of mulberry accession at the ploidy level. But, SSRs analysis is trustworthy for an understanding of the degree of genetic variation where the breeding of germplasm and mulberry germplasm conservation gets benefited. Moreover, the present research was focused to estimate genetic diversity using simple sequence repeats (SSRs), and coming to understand the phylogenetic relationship, and population structure. Incite gained herein suggested the effect of genetic variation instead of ploidy, which might be the consequence of the high level of heterozygosity imposed by natural cross-pollination.

Additionally, population structure analyses are another important aspect of the present study because it reflects the relationship between GS and marker variation in the population structure analyses. The population structure analysis revealed the existence of three genetic clusters with a high level of intermixing, a high level of within-species genetic diversity, and a weak genetic structure among *Morus* species studied. The combined clustering analysis revealed that only the majority of accessions of *M. indica* (cluster-I) and *M. macroura* (cluster-II) were grouped in a specific cluster, while the accessions of *M. alba*, *M. indica,* and *Morus* spp. were distributed throughout the three major clusters suggesting a high level of genetic admixture in these species. Such results are attributed to the outbreeding and wind pollination reproductive system of *Morus* species (Bajpai et al., 2014). In turn, the present finding suggests genetic background of species is the account of admixture between the groups that occurred due to a high level of heterozygosity as the result of gene exchange between the species during the outbreeding reproductive system. In addition, accessions were collected from different countries and they are continuously used for mulberry breeding programmes and which resulted in a mixed population. For example, *M. alba* species originating from France and Indonesia are grouped separately from those originating from Japan and India. Understanding the molecular diversity and modifications in genome content triggered by WGD, not only helps in determining the evolution but also shapes the plant genome leading to adaption, response to stress, and diversification (Soltis and Soltis, 2021). In the era of high-throughput sequencing, the genomic resources in mulberry are enriched with the chromosome-level reference genome sequencing of *M*. *alba* and draft genome sequencing of *M. indica* (Jiao et al. 2020; Jain et al. 2022) will unravel the molecular composition and genome evolution (Pellicer et al. 2018).

### Phenotypic trait plasticity and its relation to genome size (GS)

Polyploidy has been considered a source of evolutionary development, species expansion, as well as adaptation. Especially in plants, polyploids are not restricted, which occurred as a consequence of WGD events and appear to be associated with environmental conditions. Hence, the present research tried to understand the phenotypic plasticity among the ploidy groups (2x, 3x, 4x, and 6x) in *Morus* spp. However, female floral features, notably style length, were a well-recognized attribute used to classify mulberries in terms of female accessions (Minamizawa, 1976; Chang, 2006). Though, recently quantitative vegetative traits were considered and shown to be beneficial for mulberry plants (Chang et al., 2018). Moreover, in the present research, we have considered both vegetative and reproductive traits to recognize the effect of WGD on phenotypic plasticity. The correlation results indicate a significant positive correlation between GS with traits *viz*., mature inflorescence length (MIL, 0.45), and Mature fruit length (MFL, 0.38). Additionally, multiple trait-based PCA referred to GS linked with leaf area (LA), which is the account of the contribution of MIL, MFL, and InD (positive coordinate) and LLW, LLL, PT, and PL (X^+^, Y^-^ coordinate). Though, further research will require investigating how and to what extent GS is intimate to physiological and/or anatomical traits in mulberry.

## Conclusions

In the present study, we have tried to highlight the intraspecific diversity of genome size (GS), phenotypic traits, and genetic variation of a worldwide collection of 157 mulberry accessions. The primary outcome as previously mentioned fundamental queries-

1) present investigation observed that core collections bear a huge genome content variation, from diploid to hexaploids an about 4.01-fold difference of GS.

1. We have observed a significant positive correlation between GS and mature fruit length (MFL) and mature inflorescence length (MIL). The phenotypic trait plasticity indicated some trait values increased with the increased level of polyploidy. For example, leaf-lamina width and petiole width are significantly higher in tetraploids as compared to others. Correlation between GS with selected phenotypic traits was also established, however, consistency could not be defined with a higher ploidy level (>3x). In the future, to identify preferred ploidy advantages and to address ploidy-associated traits a large number of individuals with higher ploidy levels (>3x) is a prerequisite, which is restricted in the present study.
2. The population structure analysis revealed the presence of three genetic clusters with a high level of admixture, a weak genetic structure, and a high level of genetic diversity within *Morus* spp., which implies the effect of genetic variation instead of ploidy on trait plasticity that could be a consequence of the high level of heterozygosity imposed by natural cross-pollination. Polyploidy in mulberry may be due to several different sources of origin. In general, polyploids have considerable value in crop plants, particularly crops that are grown for their vegetative parts and they play an important role in the natural selection and better adaptability of species in new ecological niches. Identification of these polyploids through cytomorphological analysis is quite difficult in mulberry as their huge variation in chromosome size and number. Hence, the present FCM analysis was useful in distinguishing the mulberry at different ploidy levels, which provides information about the variation level of GC among the different ploidy levels of mulberry germplasm for future research and conservation.

## Acknowledgments

Dr. Gnanesh B.N. acknowledges SERB, for providing Ramanujan Fellowship. The financial support was provided by Science and Engineering Research Board (SERB), New Delhi, grant (SERB SB/S2/RJN-049/2015). Mr. Raju Mondal acknowledged cytogenetics work supported by Central Silk Board (CSB), Bangalore, grant (PIG06004 SI). Thanks to Karnataka State Sericulture Research and Development Institute (KSSRDI), Bangalore, Karnataka, India for providing the germplasm materials.

## Author’s contributions

GBN – Conceive the idea, fund acquisition, and designed the study, RM-Cytology work, morphological data analysis, interpretation, and presentation; GBN & RM - original draft preparation and final editing of the manuscript with the help of other authors; GBN, MHB &SMB-Genome content analysis; MHB, PS & BNG-performed the molecular experiments, data analysis, and interpretation; BMR, BNG, SP & AGS, - morphological data collection and analysis; MT, SV-Provided the essential genotypes and proof editing; All authors read and approved the manuscript

## Funding

This research was funded by Science and Engineering Research Board (SERB), New Delhi, grant number SERB SB/S2/RJN-049/2015.

## Competing Interests

The authors declare there are no conflicts of interests

## Ethical approval

This article does not contain any studies with human or animal participants

## Notes

### Competing Interest Statement

The authors have declared no competing interest.

