## Supplementary material for "Genome size, genetic diversity, and phenotypic variability imply the effect of genetic variation instead of ploidy on trait plasticity in the cross-pollinated tree species of mulberry": S.Fig 2

### Slide 1
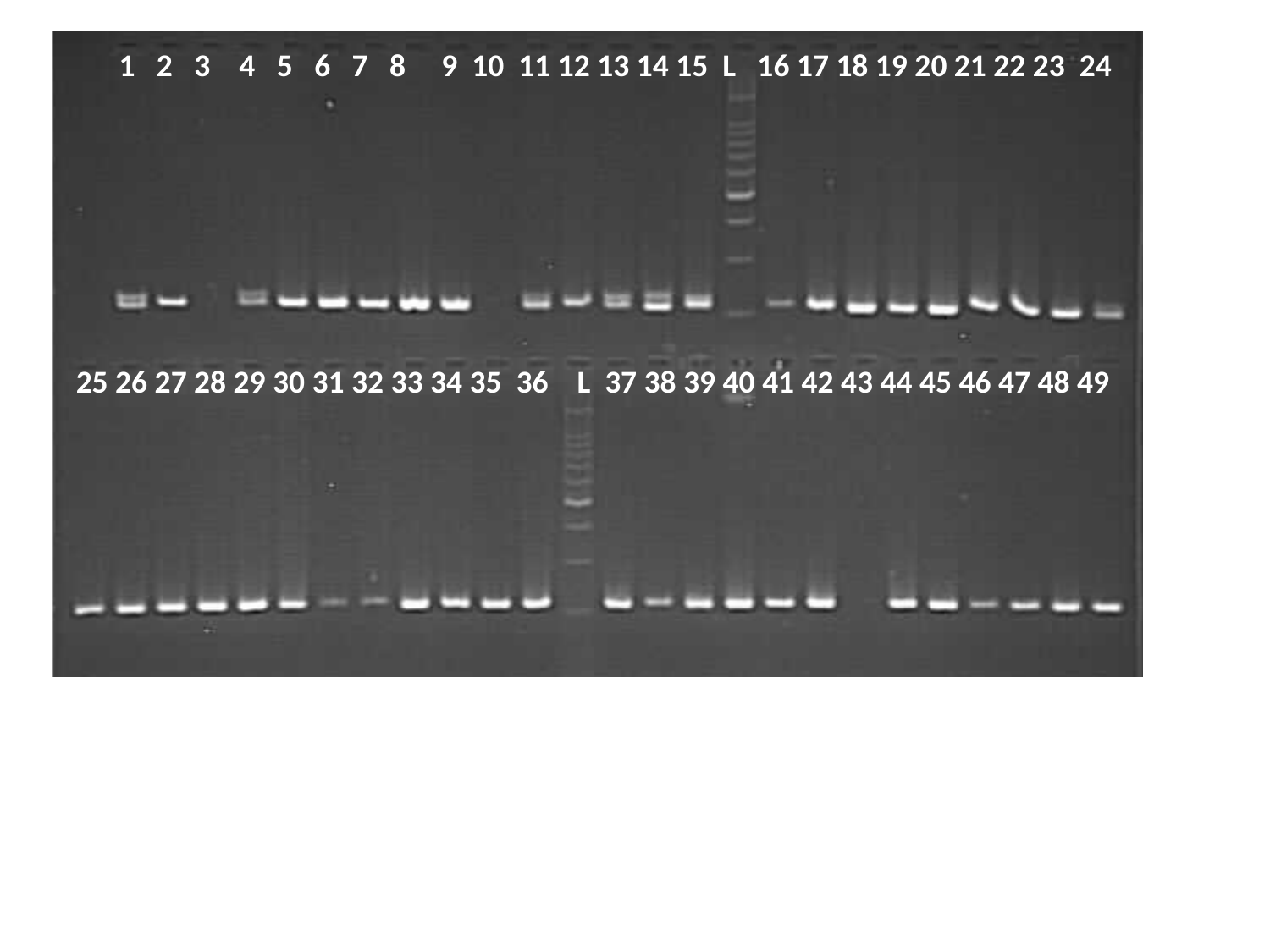

1 2 3 4 5 6 7 8 9 10 11 12 13 14 15 L 16 17 18 19 20 21 22 23 24
25 26 27 28 29 30 31 32 33 34 35 36 L 37 38 39 40 41 42 43 44 45 46 47 48 49

### Slide 2
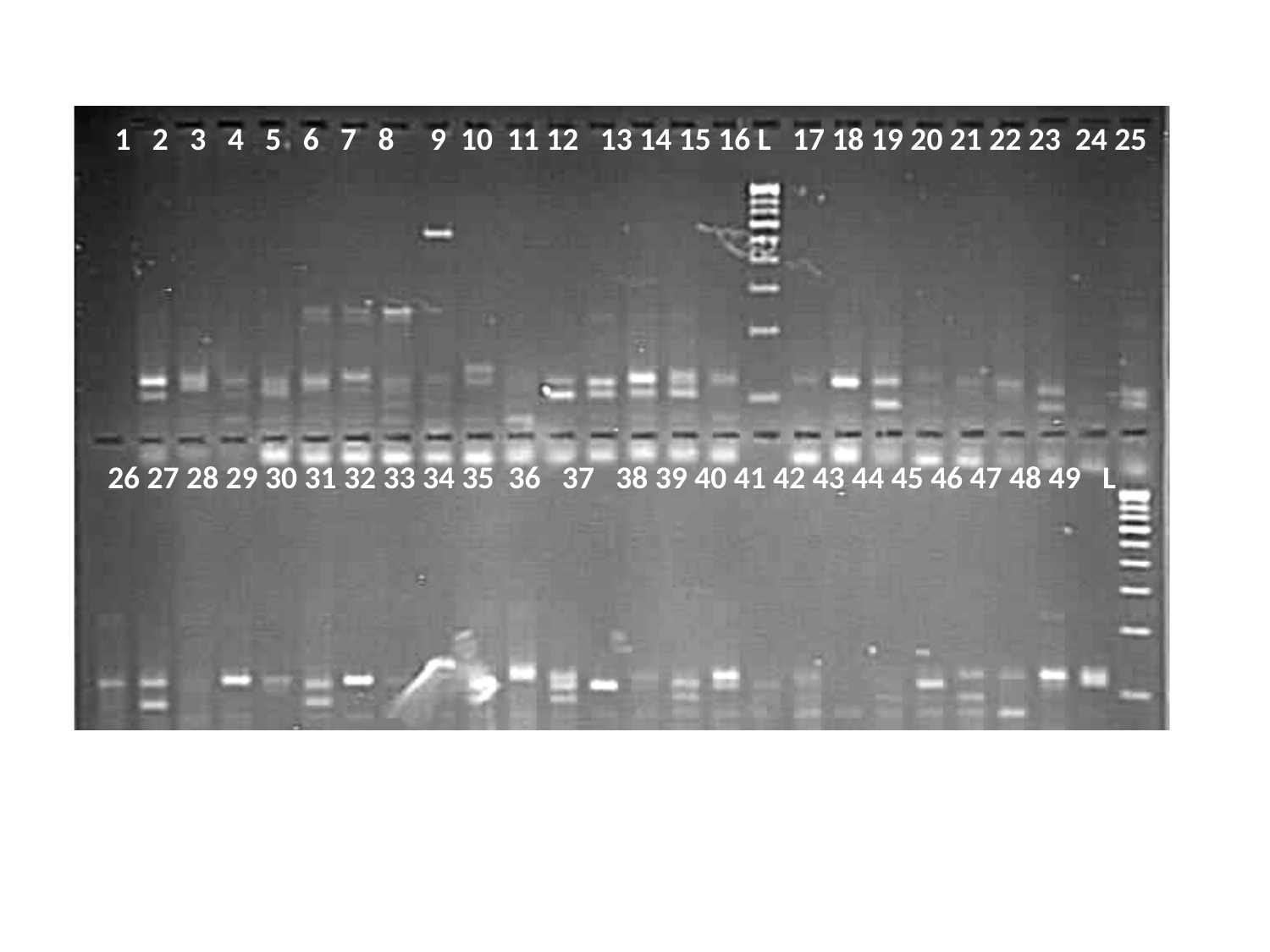

1 2 3 4 5 6 7 8 9 10 11 12 13 14 15 16 L 17 18 19 20 21 22 23 24 25
26 27 28 29 30 31 32 33 34 35 36 37 38 39 40 41 42 43 44 45 46 47 48 49 L

### Slide 3
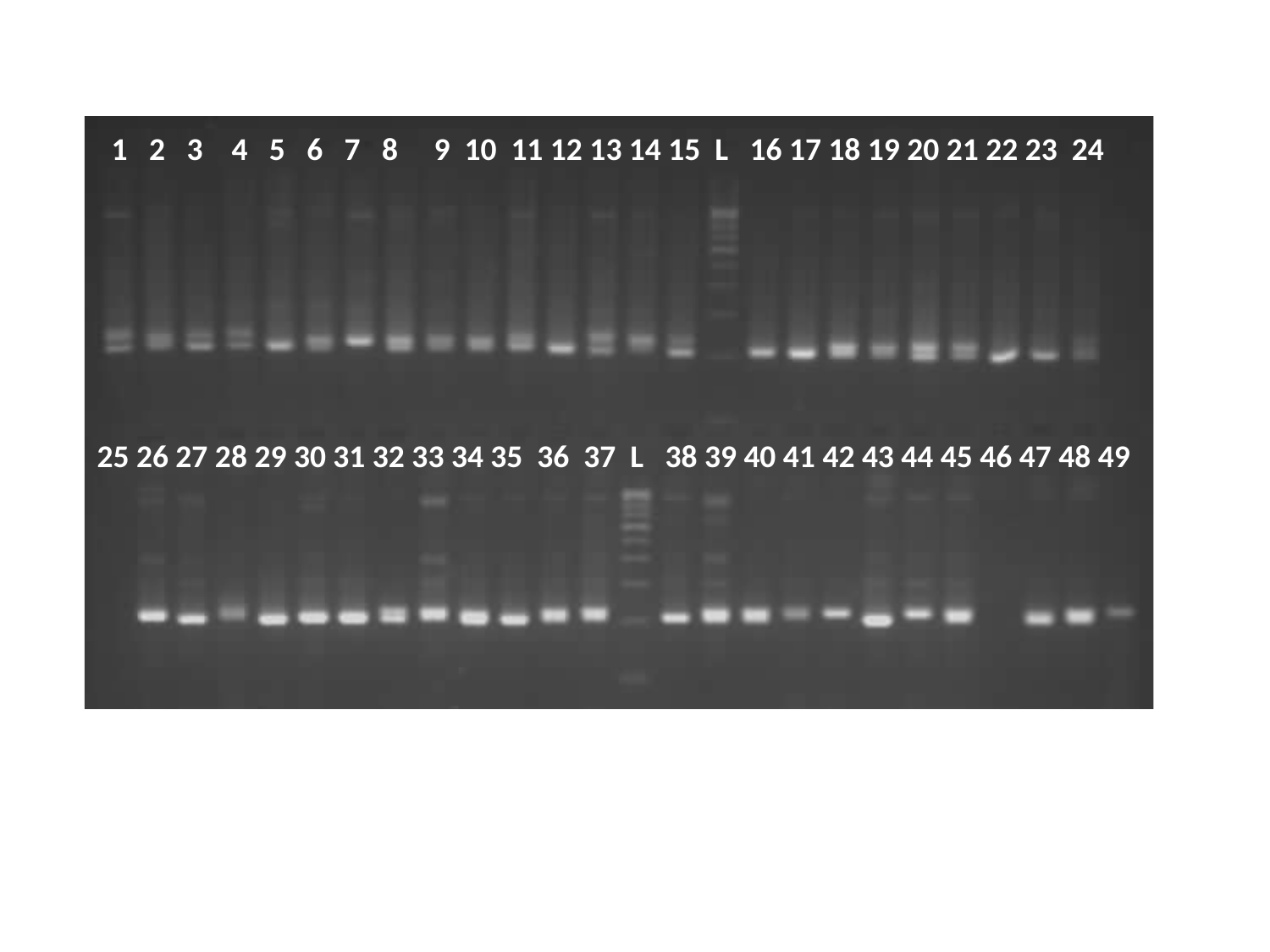

1 2 3 4 5 6 7 8 9 10 11 12 13 14 15 L 16 17 18 19 20 21 22 23 24
25 26 27 28 29 30 31 32 33 34 35 36 37 L 38 39 40 41 42 43 44 45 46 47 48 49

### Slide 4
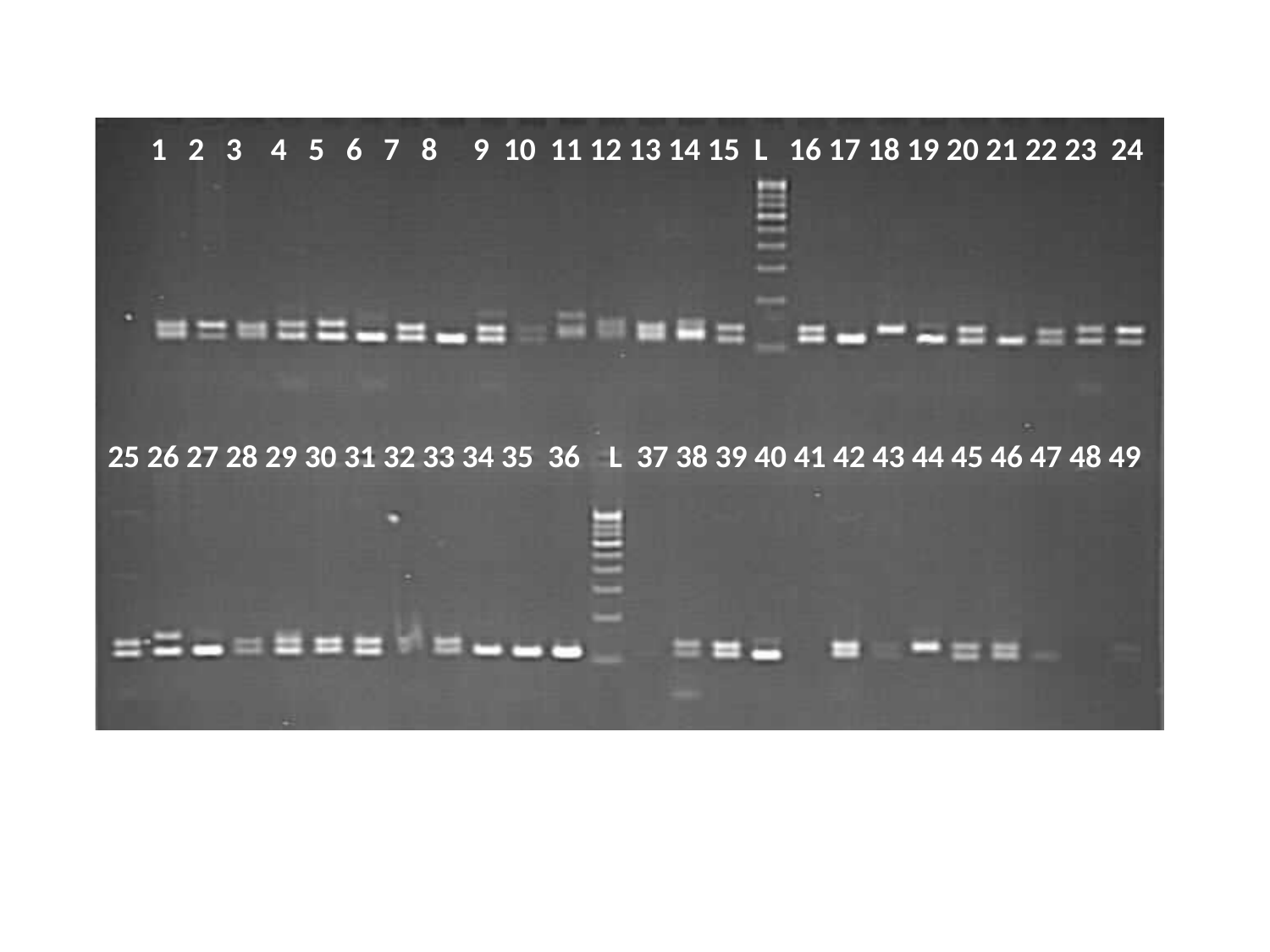

1 2 3 4 5 6 7 8 9 10 11 12 13 14 15 L 16 17 18 19 20 21 22 23 24
25 26 27 28 29 30 31 32 33 34 35 36 L 37 38 39 40 41 42 43 44 45 46 47 48 49

### Slide 5
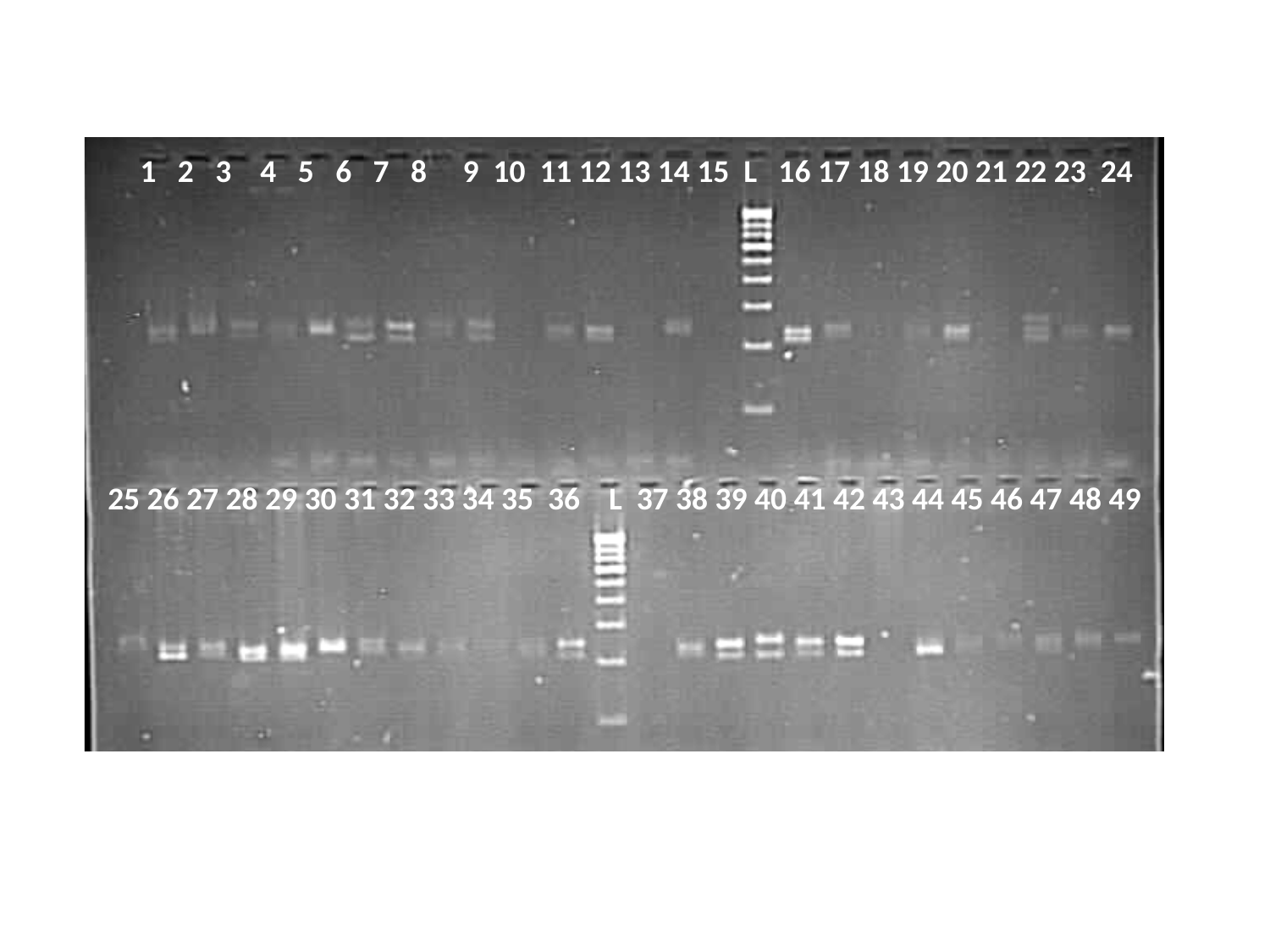

1 2 3 4 5 6 7 8 9 10 11 12 13 14 15 L 16 17 18 19 20 21 22 23 24
25 26 27 28 29 30 31 32 33 34 35 36 L 37 38 39 40 41 42 43 44 45 46 47 48 49

### Slide 6
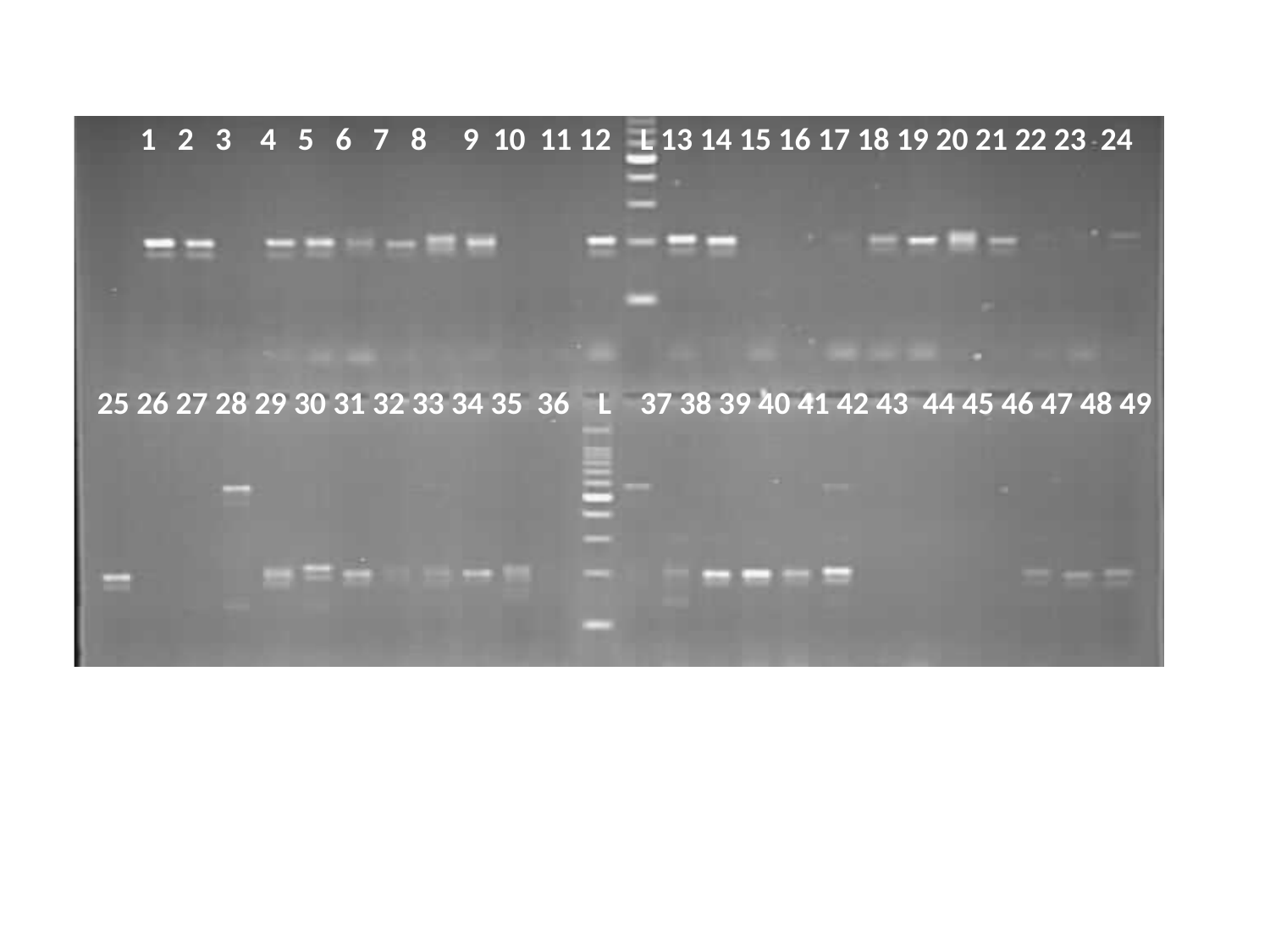

1 2 3 4 5 6 7 8 9 10 11 12 L 13 14 15 16 17 18 19 20 21 22 23 24
25 26 27 28 29 30 31 32 33 34 35 36 L 37 38 39 40 41 42 43 44 45 46 47 48 49

### Slide 7
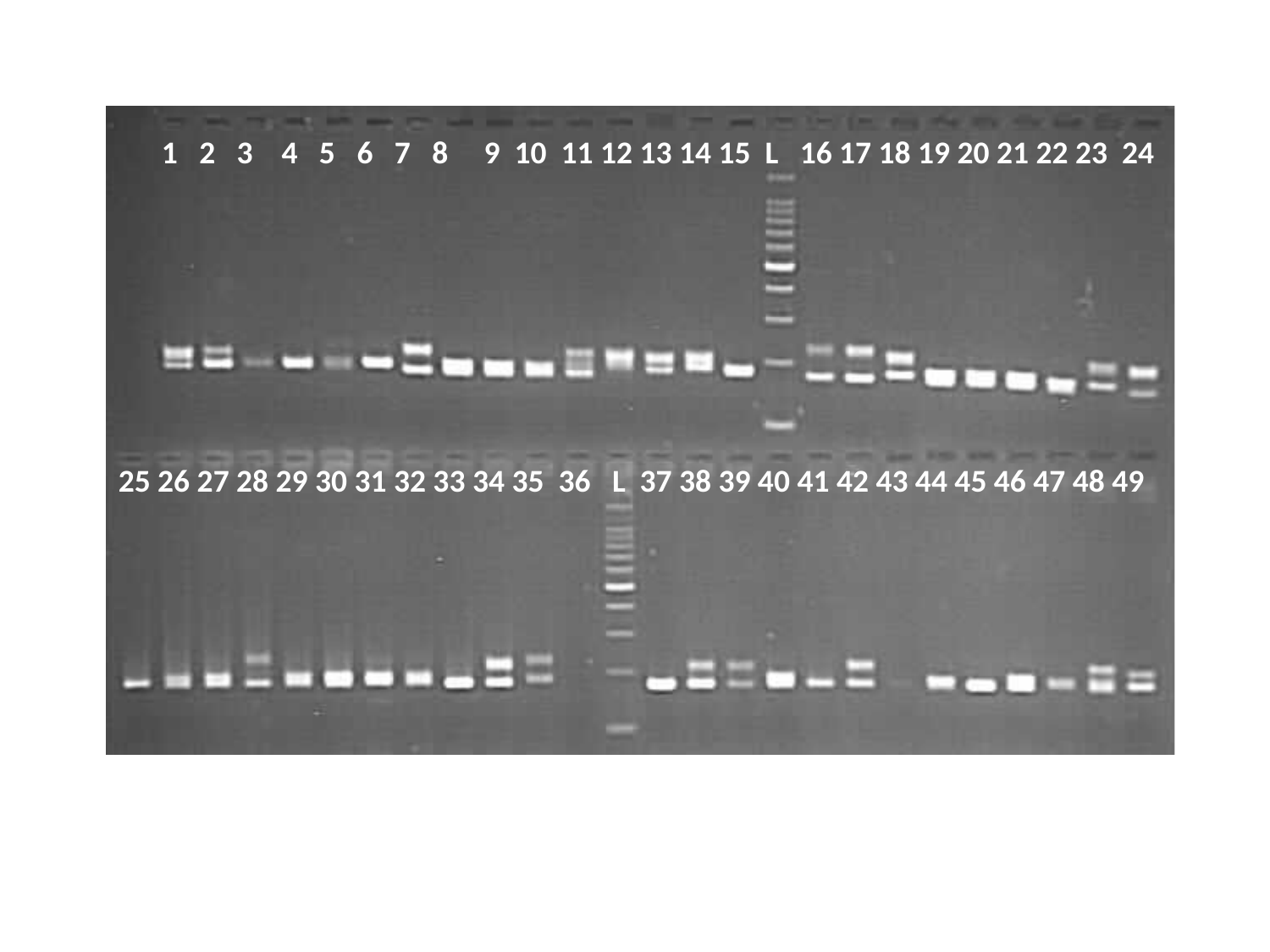

1 2 3 4 5 6 7 8 9 10 11 12 13 14 15 L 16 17 18 19 20 21 22 23 24
25 26 27 28 29 30 31 32 33 34 35 36 L 37 38 39 40 41 42 43 44 45 46 47 48 49

### Slide 8
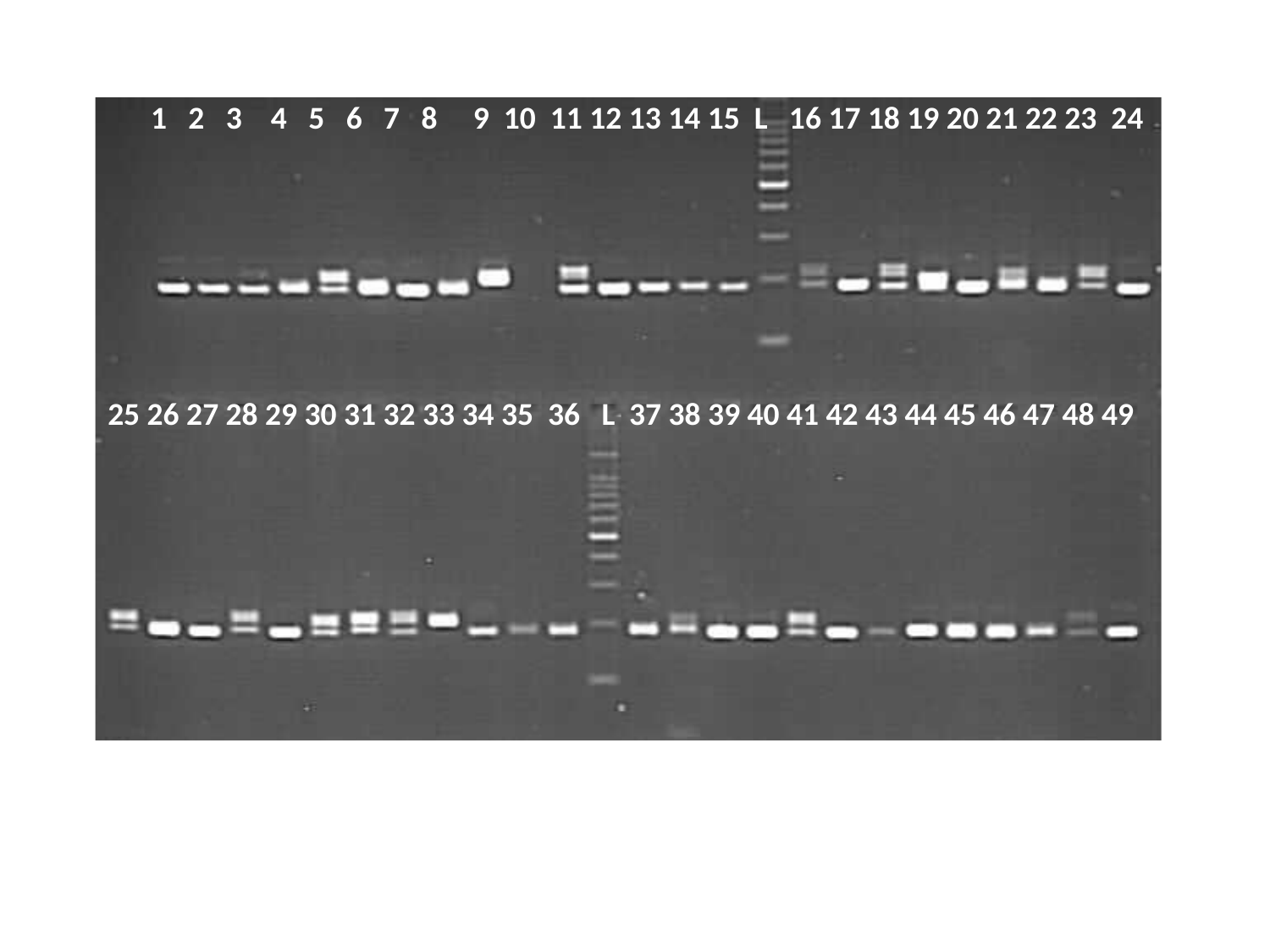

1 2 3 4 5 6 7 8 9 10 11 12 13 14 15 L 16 17 18 19 20 21 22 23 24
25 26 27 28 29 30 31 32 33 34 35 36 L 37 38 39 40 41 42 43 44 45 46 47 48 49

### Slide 9
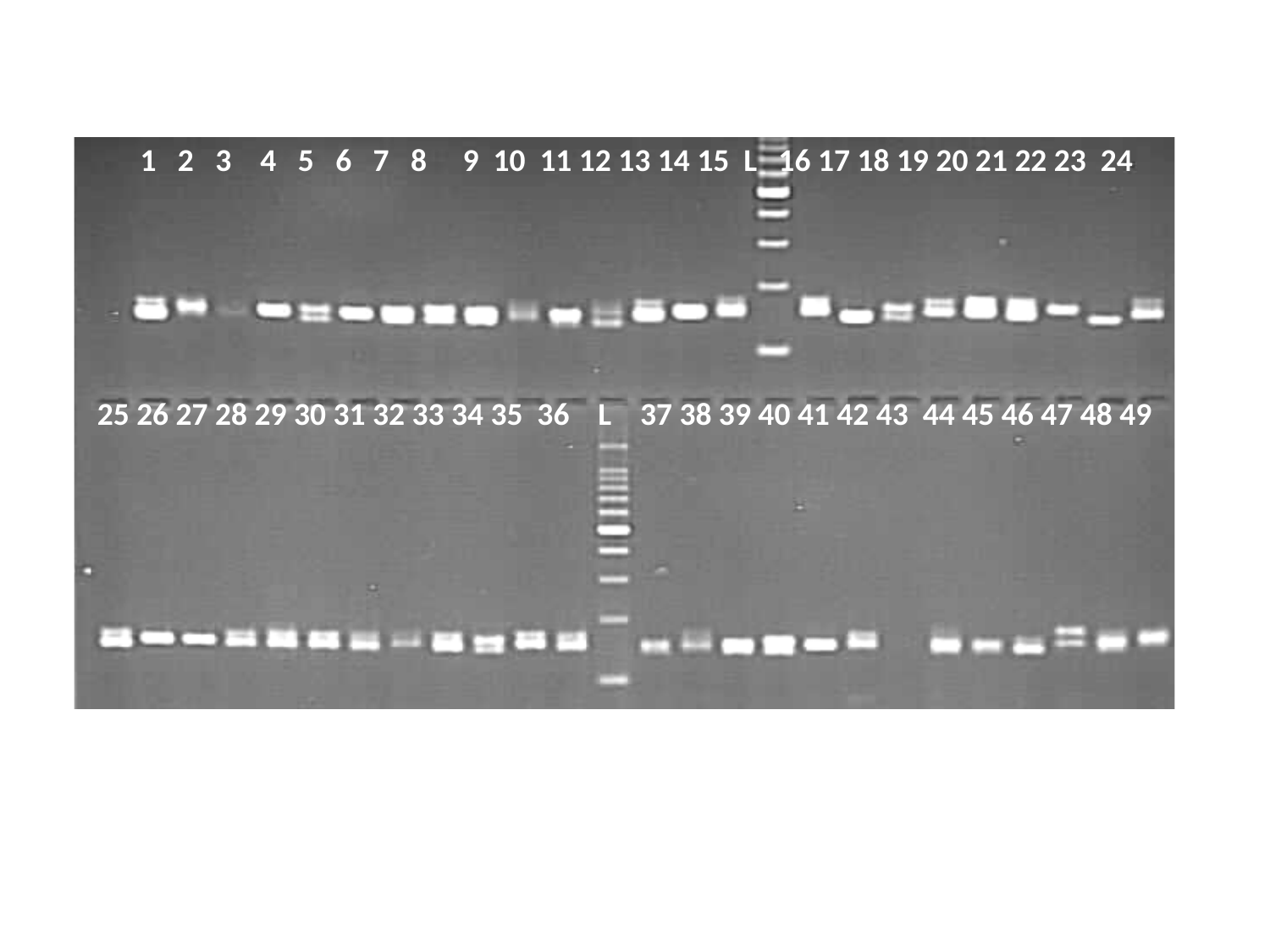

1 2 3 4 5 6 7 8 9 10 11 12 13 14 15 L 16 17 18 19 20 21 22 23 24
25 26 27 28 29 30 31 32 33 34 35 36 L 37 38 39 40 41 42 43 44 45 46 47 48 49

### Slide 10
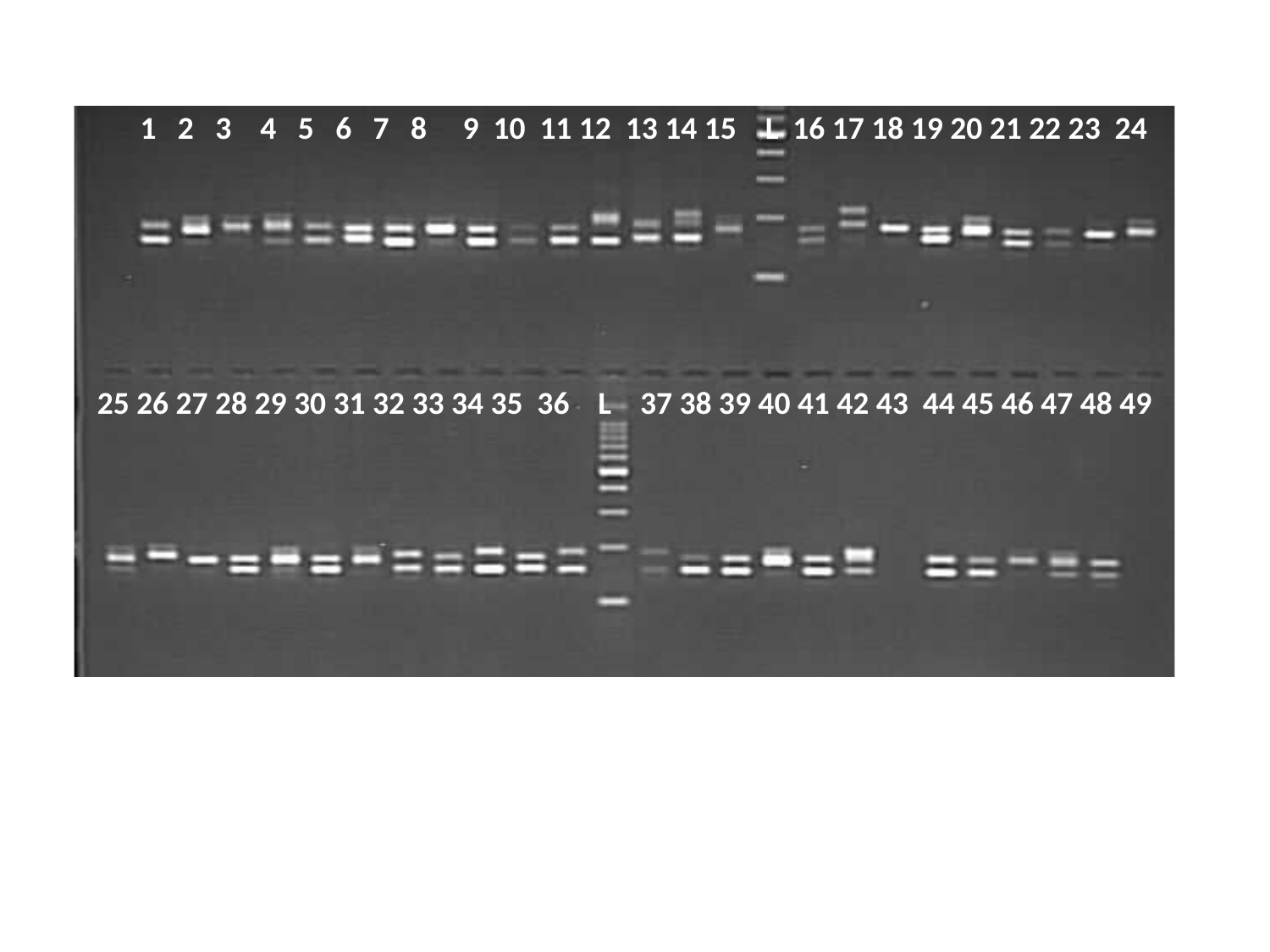

1 2 3 4 5 6 7 8 9 10 11 12 13 14 15 L 16 17 18 19 20 21 22 23 24
25 26 27 28 29 30 31 32 33 34 35 36 L 37 38 39 40 41 42 43 44 45 46 47 48 49

### Slide 11
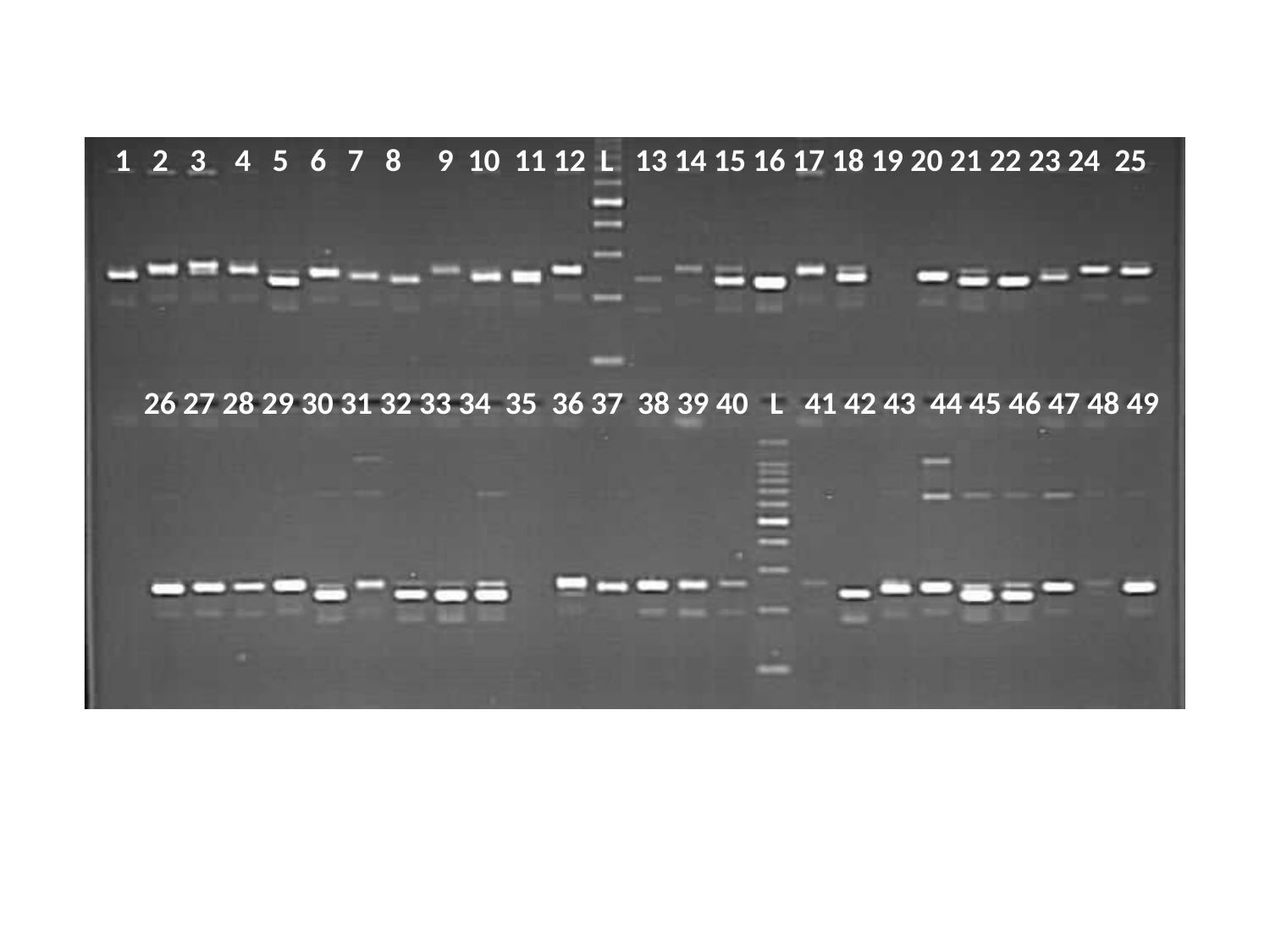

1 2 3 4 5 6 7 8 9 10 11 12 L 13 14 15 16 17 18 19 20 21 22 23 24 25
 26 27 28 29 30 31 32 33 34 35 36 37 38 39 40 L 41 42 43 44 45 46 47 48 49

### Slide 12
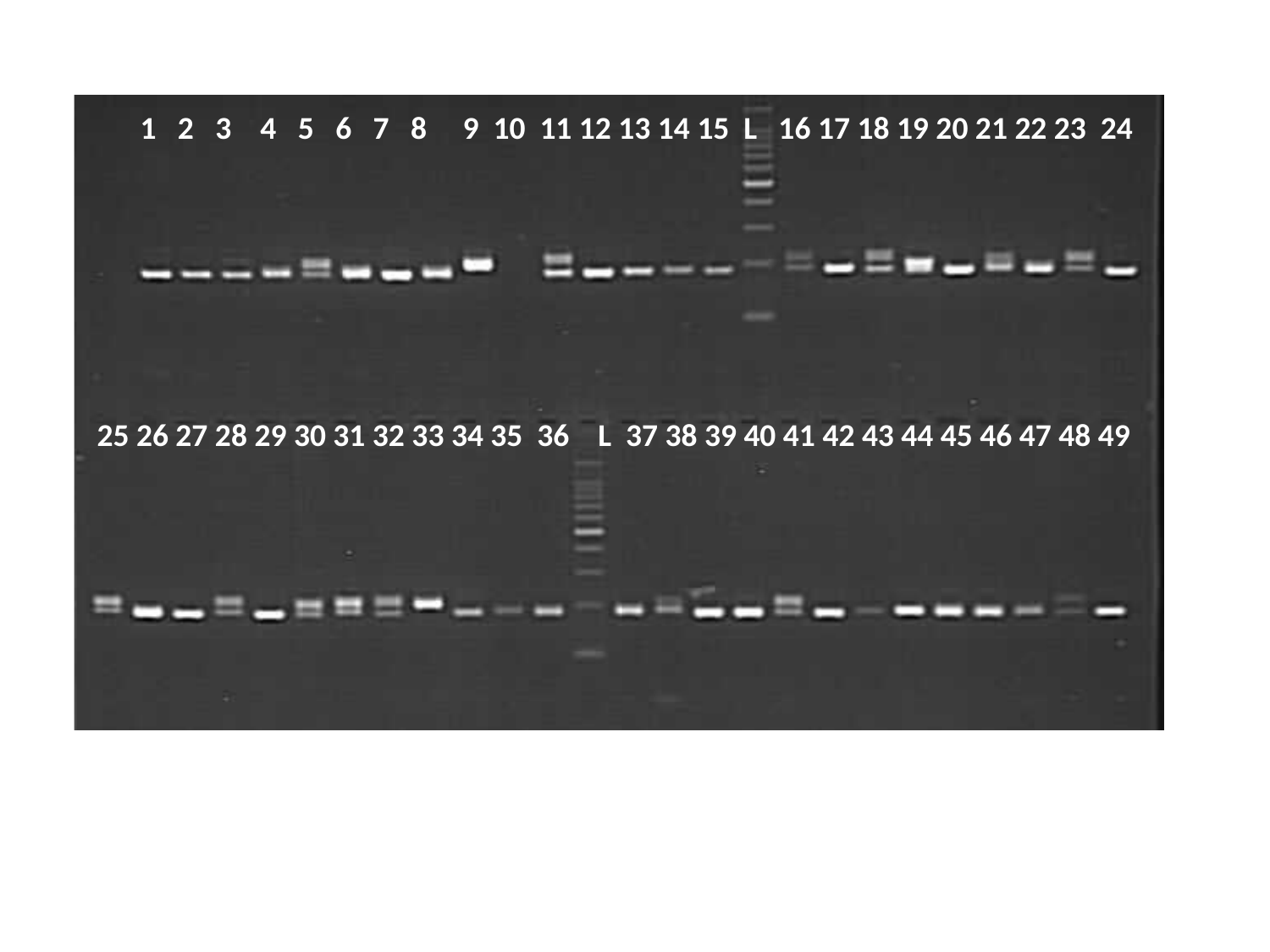

1 2 3 4 5 6 7 8 9 10 11 12 13 14 15 L 16 17 18 19 20 21 22 23 24
25 26 27 28 29 30 31 32 33 34 35 36 L 37 38 39 40 41 42 43 44 45 46 47 48 49

### Slide 13
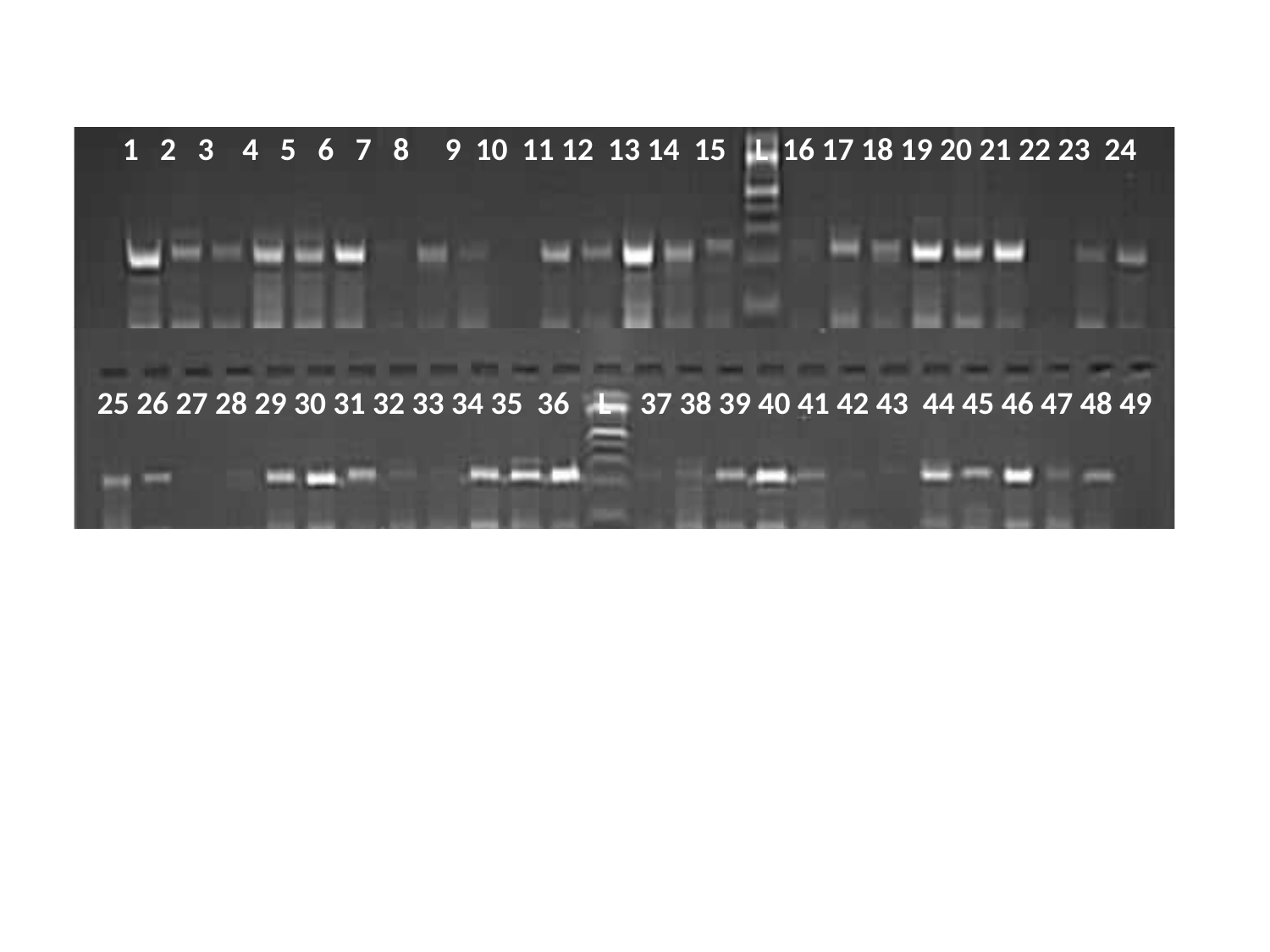

1 2 3 4 5 6 7 8 9 10 11 12 13 14 15 L 16 17 18 19 20 21 22 23 24
25 26 27 28 29 30 31 32 33 34 35 36 L 37 38 39 40 41 42 43 44 45 46 47 48 49

### Slide 14
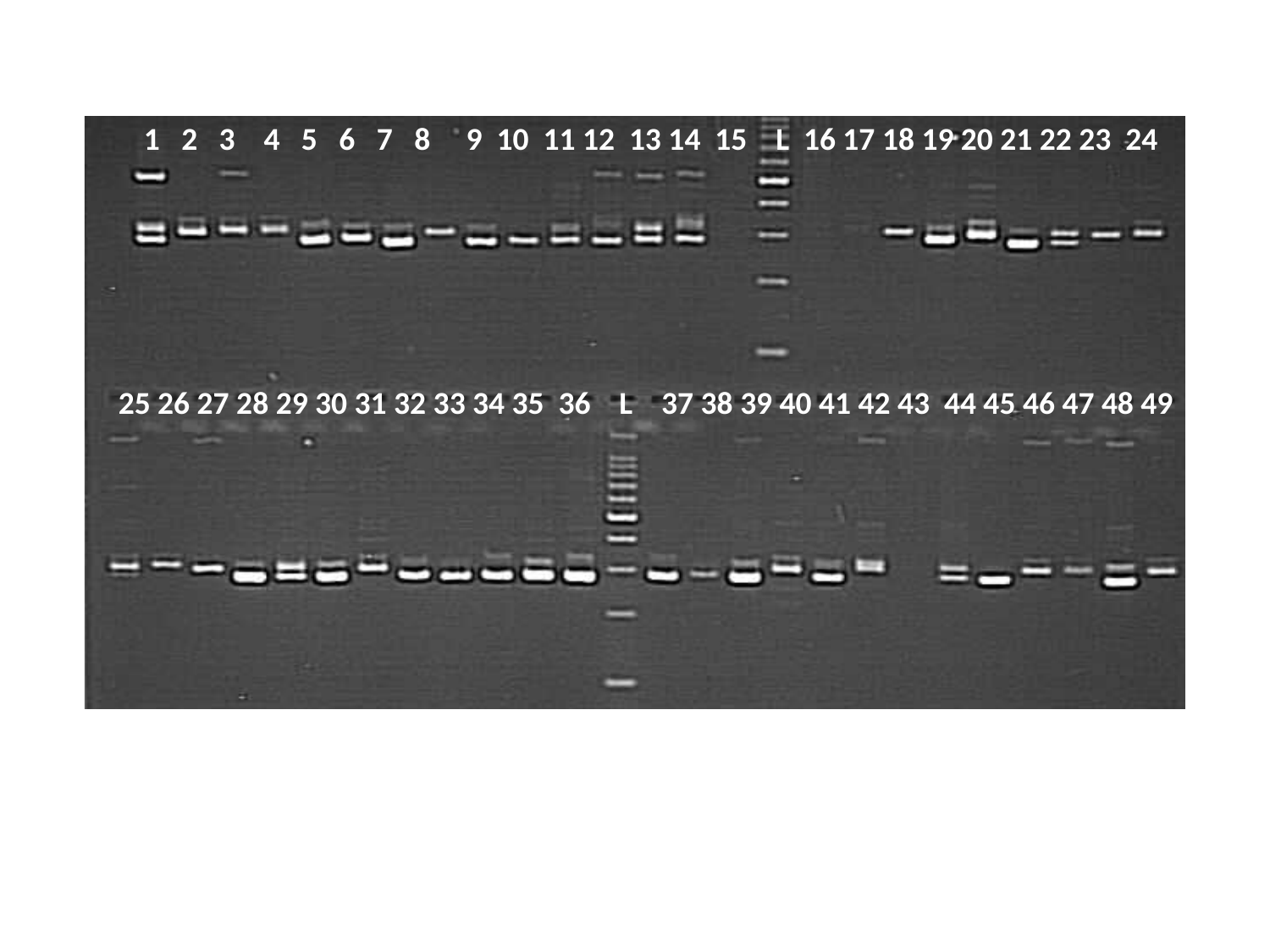

1 2 3 4 5 6 7 8 9 10 11 12 13 14 15 L 16 17 18 19 20 21 22 23 24
25 26 27 28 29 30 31 32 33 34 35 36 L 37 38 39 40 41 42 43 44 45 46 47 48 49

### Slide 15
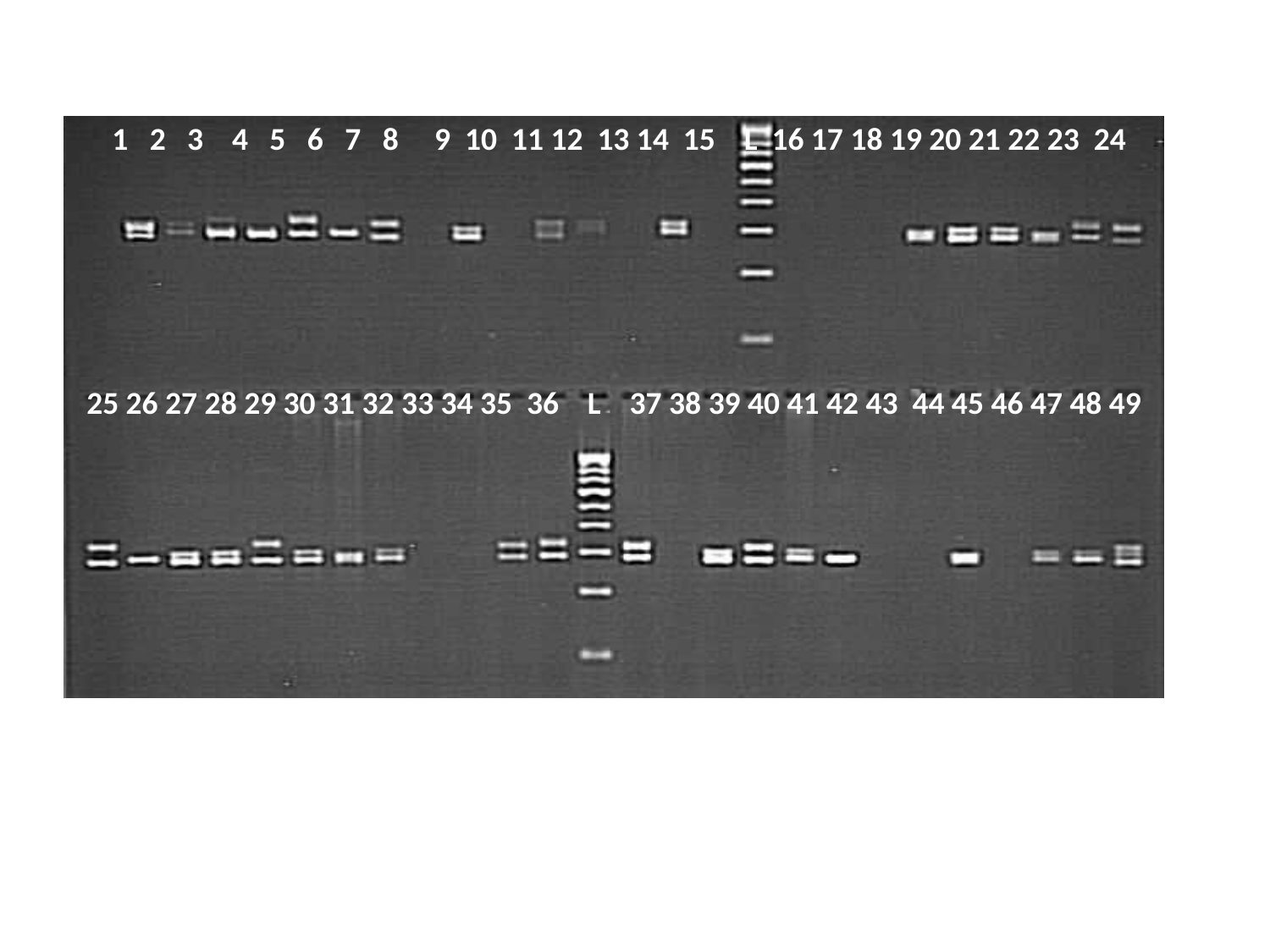

1 2 3 4 5 6 7 8 9 10 11 12 13 14 15 L 16 17 18 19 20 21 22 23 24
25 26 27 28 29 30 31 32 33 34 35 36 L 37 38 39 40 41 42 43 44 45 46 47 48 49

### Slide 16
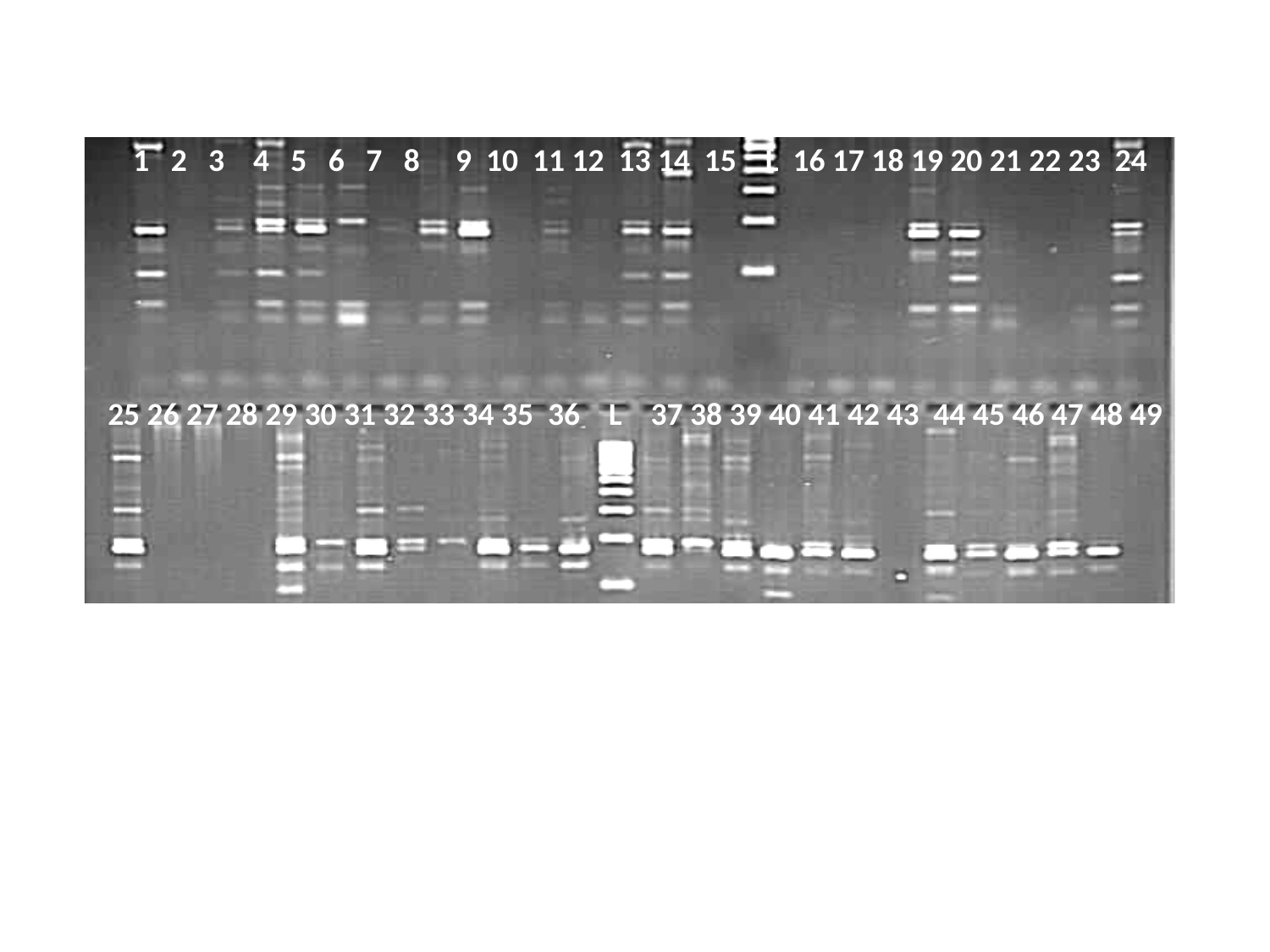

1 2 3 4 5 6 7 8 9 10 11 12 13 14 15 L 16 17 18 19 20 21 22 23 24
25 26 27 28 29 30 31 32 33 34 35 36 L 37 38 39 40 41 42 43 44 45 46 47 48 49

### Slide 17
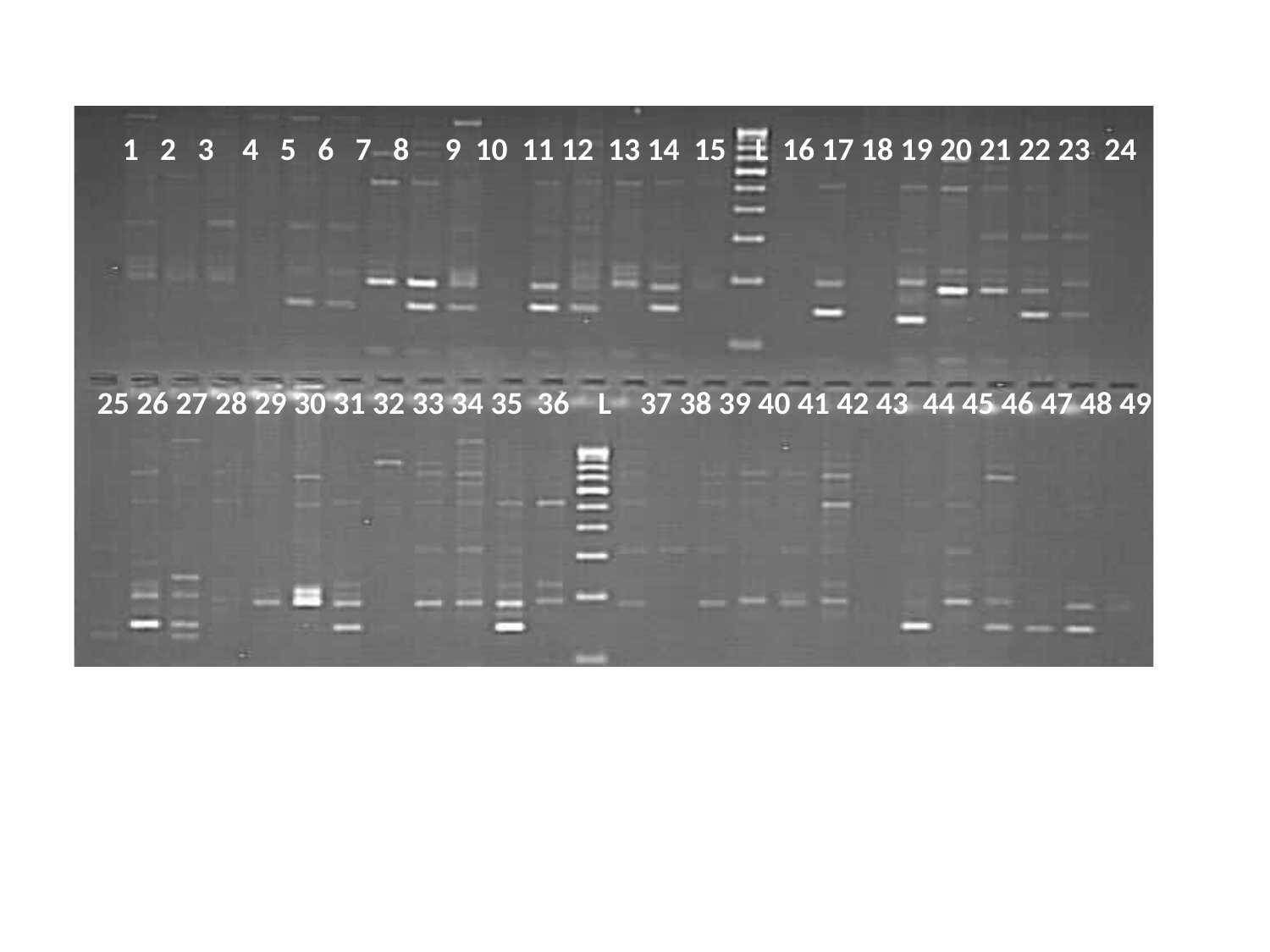

1 2 3 4 5 6 7 8 9 10 11 12 13 14 15 L 16 17 18 19 20 21 22 23 24
25 26 27 28 29 30 31 32 33 34 35 36 L 37 38 39 40 41 42 43 44 45 46 47 48 49

### Slide 18
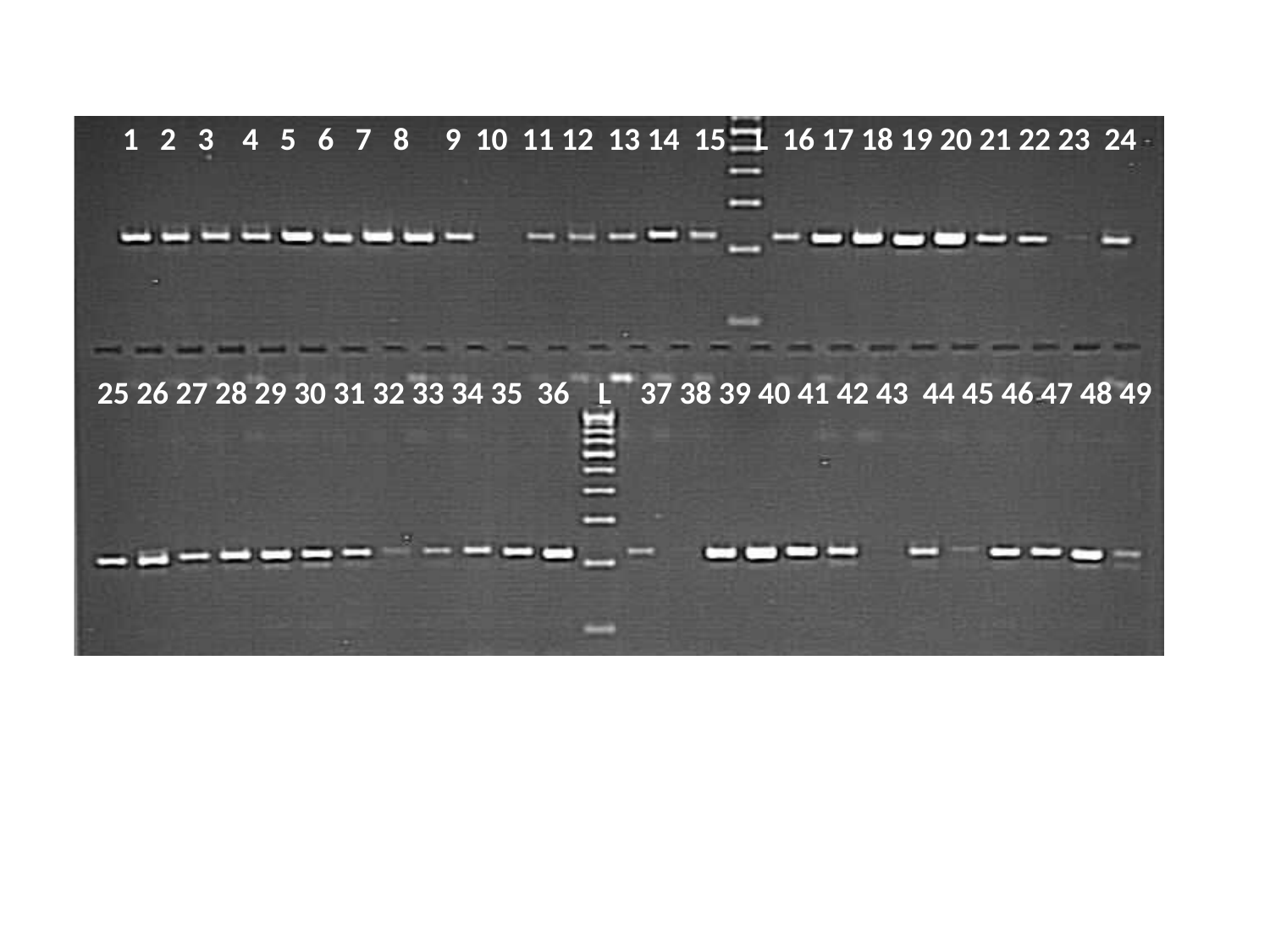

1 2 3 4 5 6 7 8 9 10 11 12 13 14 15 L 16 17 18 19 20 21 22 23 24
25 26 27 28 29 30 31 32 33 34 35 36 L 37 38 39 40 41 42 43 44 45 46 47 48 49

### Slide 19
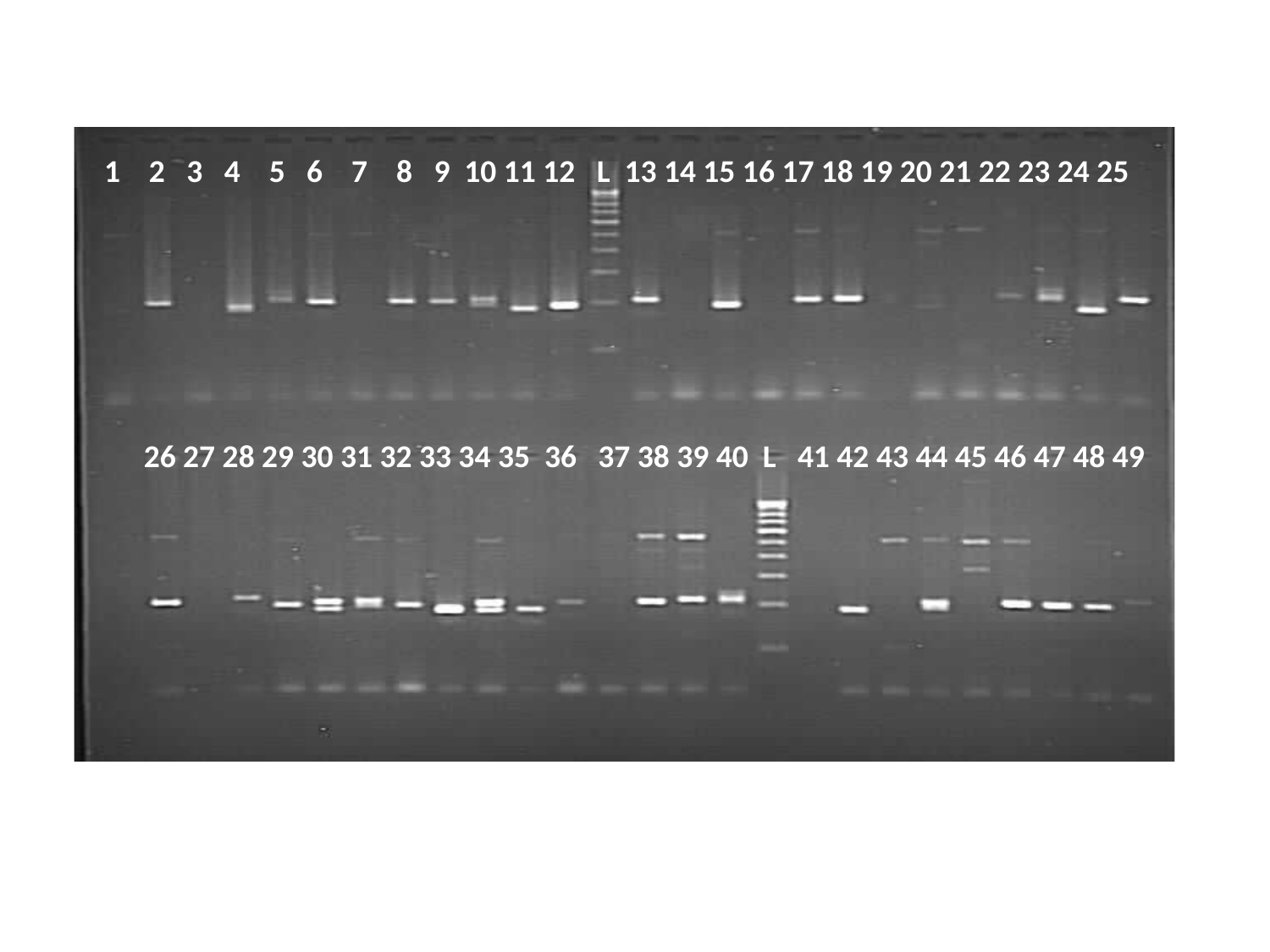

1 2 3 4 5 6 7 8 9 10 11 12 L 13 14 15 16 17 18 19 20 21 22 23 24 25
 26 27 28 29 30 31 32 33 34 35 36 37 38 39 40 L 41 42 43 44 45 46 47 48 49

### Slide 20
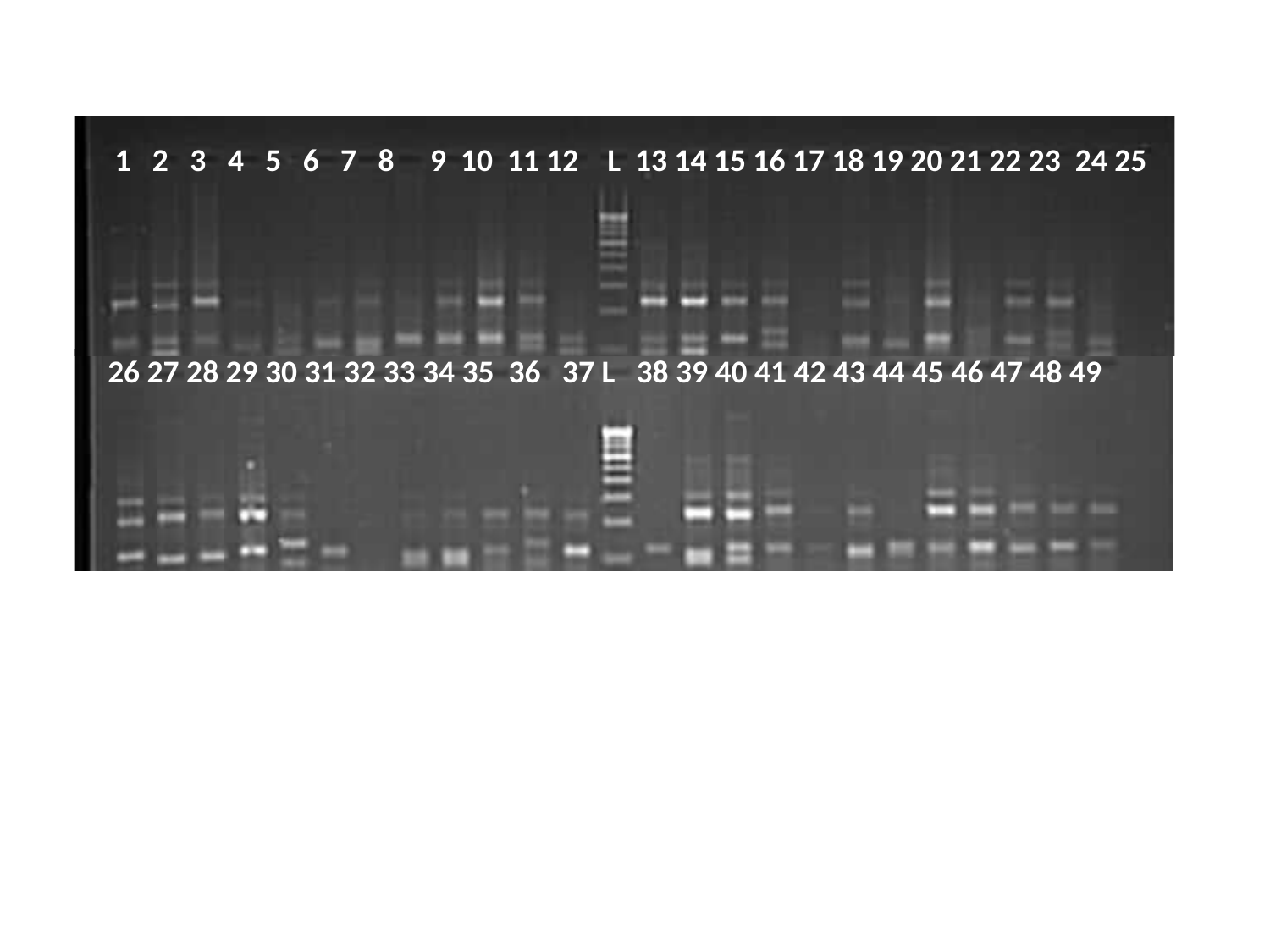

1 2 3 4 5 6 7 8 9 10 11 12 L 13 14 15 16 17 18 19 20 21 22 23 24 25
26 27 28 29 30 31 32 33 34 35 36 37 L 38 39 40 41 42 43 44 45 46 47 48 49
