## Supplementary material for "Genome size, genetic diversity, and phenotypic variability imply the effect of genetic variation instead of ploidy on trait plasticity in the cross-pollinated tree species of mulberry": S.Fig 1

**Figure S1** (A) Origin and distribution of worldwide mulberry collections used in this study. (B) The donut chart indicates the proportion of genotype collections corresponding to the country.

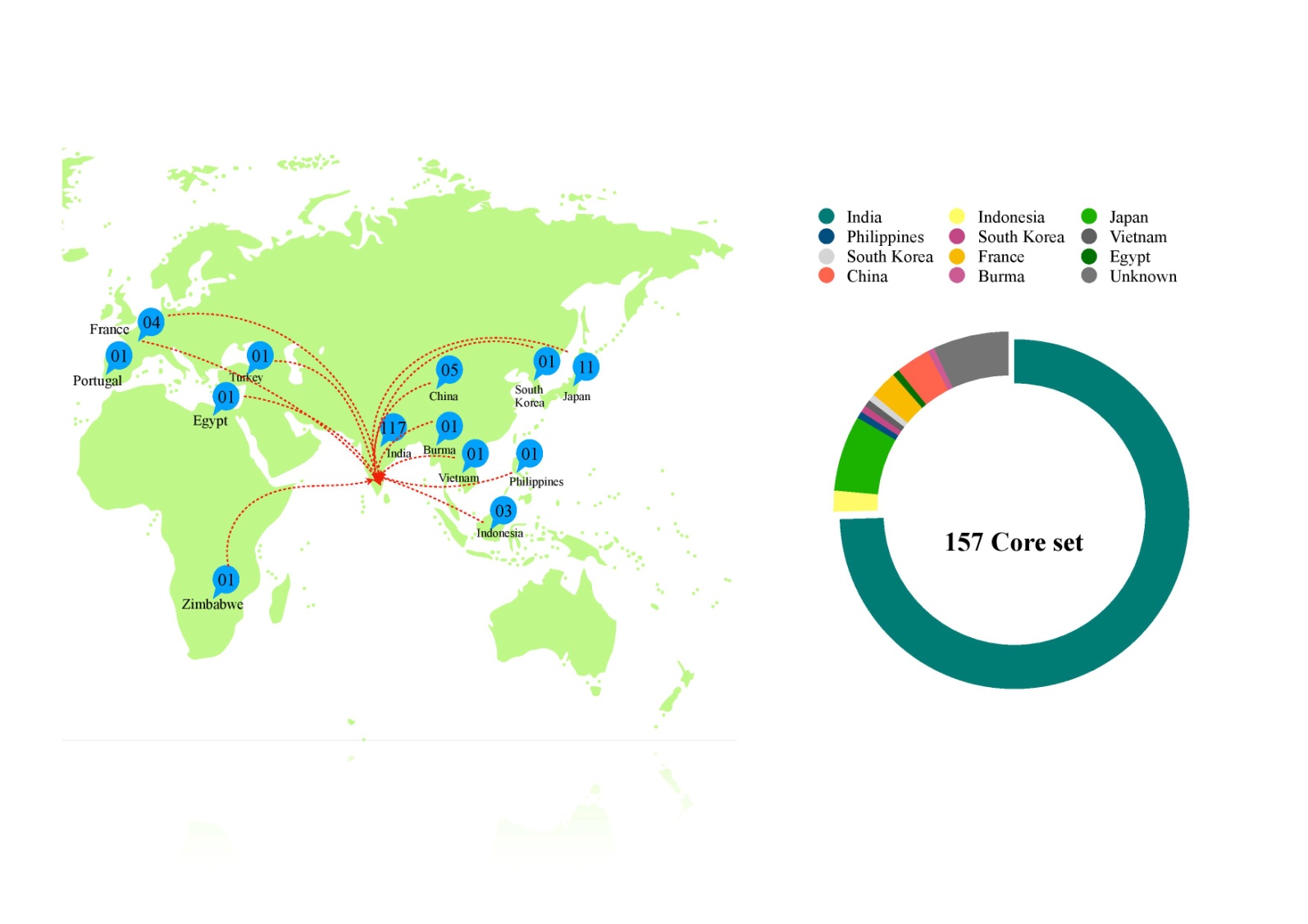
